# Resting-state functional brain connectivity best predicts the personality dimension of openness to experience

**DOI:** 10.1101/215129

**Authors:** Julien Dubois, Paola Galdi, Yanting Han, Lynn K. Paul, Ralph Adolphs

## Abstract

Personality neuroscience aims to find associations between brain measures and personality traits. Findings to date have been severely limited by a number of factors, including small sample size and omission of out-of-sample prediction. We capitalized on the recent availability of a large database, together with the emergence of specific criteria for best practices in neuroimaging studies of individual differences. We analyzed resting-state functional magnetic resonance imaging data from 884 young healthy adults in the Human Connectome Project (HCP) database. We attempted to predict personality traits from the “Big Five”, as assessed with the NEO-FFI test, using individual functional connectivity matrices. After regressing out potential confounds (such as age, sex, handedness and fluid intelligence), we used a cross-validated framework, together with test-retest replication (across two sessions of resting-state fMRI for each subject), to quantify how well the neuroimaging data could predict each of the five personality factors. We tested three different (published) denoising strategies for the fMRI data, two inter-subject alignment and brain parcellation schemes, and three different linear models for prediction. As measurement noise is known to moderate statistical relationships, we performed final prediction analyses using average connectivity across both imaging sessions (1 h of data), with the analysis pipeline that yielded the highest predictability overall. Across all results (test/retest; 3 denoising strategies; 2 alignment schemes; 3 models), Openness to experience emerged as the only reliably predicted personality factor. Using the full hour of resting-state data and the best pipeline, we could predict Openness to experience (NEOFAC_O: r=0.24, R^2^=0.024) almost as well as we could predict the score on a 24-item intelligence test (PMAT24_A_CR: r=0.26, R^2^=0.044). Other factors (Extraversion, Neuroticism, Agreeableness and Conscientiousness) yielded weaker predictions across results that were not statistically significant under permutation testing. We also derived two superordinate personality factors (“α” and “β”) from a principal components analysis of the NEO-FFI factor scores, thereby reducing noise and enhancing the precision of these measures of personality. We could account for 5% of the variance in the β superordinate factor (r=0.27, R^2^=0.050), which loads highly on Openness to experience. We conclude with a discussion of the potential for predicting personality from neuroimaging data and make specific recommendations for the field.

## Introduction

Personality refers to the relatively stable disposition of an individual that influences long-term behavioral style (Back, Schmukle, & Egloff, 2009; Furr, 2009; Hong, Paunonen, & Slade, 2008; Jaccard, 1974). It is especially conspicuous in social interactions, and in emotional expression. It is what we pick up on when we observe a person for an extended time, and what leads us to make predictions about general tendencies in behaviors and interactions in the future. Often, these predictions are inaccurate stereotypes, and they can be evoked even by very fleeting impressions, such as merely looking at photographs of people (Todorov, 2017). Yet there is also good reliability (Viswesvaran & Ones, 2000) and consistency (B. W. Roberts & DelVecchio, 2000) for many personality traits currently used in psychology, which can predict real-life outcomes (Brent W. Roberts, Kuncel, Shiner, Caspi, & Goldberg, 2007).

While human personality traits are typically inferred from questionnaires, viewed as latent variables they could plausibly be derived also from other measures. In fact, there are good reasons to think that biological measures other than self-reported questionnaires can be used to estimate personality traits. Many of the personality traits similar to those used to describe human dispositions can be applied to animal behavior as well, and again they make some predictions about real-life outcomes (Gosling & John, 1999; Gosling & Vazire, 2002). For instance, anxious temperament has been a major topic of study in monkeys, as a model of human mood disorders. Hyenas show neuroticism in their behavior, and also show sex differences in this trait as would be expected from human data (in humans, females tend to be more neurotic than males; in hyenas, the females are socially dominant and the males are more neurotic). Personality traits are also highly heritable. Anxious temperament in monkeys is heritable and its neurobiological basis is being intensively investigated (Oler et al., 2010). Twin studies in humans typically report heritability estimates for each trait between .4 and .6 (Bouchard & McGue, 2003; Jang, Livesley, & Vernon, 1996; Verweij et al., 2010), even though no individual genes account for much variance (studies using common single-nucleotide polymorphisms (SNPs) report estimates between 0 and 0.2 (R. A. Power & Pluess, 2015; Vinkhuyzen et al., 2012)).

Just as gene-environment interactions constitute the distal causes of our phenotype, the proximal cause of personality must come from brain-environment interactions, since these are the basis for all behavioral patterns. Some aspects of personality have been linked to specific neural systems -- for instance, behavioral inhibition and anxious temperament have been linked to a system involving the medial temporal lobe and the prefrontal cortex (Birn et al., 2014). Although there is now universal agreement that personality is generated through brain function in a given context, it is much less clear what type of brain measure might be the best predictor of personality. Neurotransmitters, cortical thickness or volume of certain regions, and functional measures have all been explored with respect to their correlation with personality traits (see (Turhan Canli, 2006; Yarkoni., 2015) for reviews). We briefly summarize this literature next and refer the interested reader to review articles and primary literature for the details.

### The search for neurobiological substrates of personality traits

Since personality traits are relatively stable over time (unlike state variables, such as emotions), one might expect that brain measures that are similarly stable over time are the most promising candidates for predicting such traits. The first types of measures to look at might thus be structural, connectional and neurochemical; indeed a number of such studies have reported correlations with personality differences. Here we briefly review studies using structural and functional magnetic resonance imaging (MRI) of humans, but leave aside research on neurotransmission (see (Turhan Canli, 2006; Yarkoni., 2015) for more exhaustive reviews). Although a number of different personality traits have been investigated, we emphasize those most similar to the “Big Five”, since they are the topic of the present paper (see below).

#### Structural MRI studies

Many structural MRI studies of personality to date have used voxel-based morphometry (VBM) (Blankstein, Chen, Mincic, McGrath, & Davis, 2009; Coutinho, Sampaio, Ferreira, Soares, & Gonçalves, 2013; DeYoung et al., 2010; Hu et al., 2011; Kapogiannis, Sutin, Davatzikos, Costa, & Resnick, 2013; W.-Y. Liu et al., 2013; Lu et al., 2014; Omura, Todd Constable, & Canli, 2005; Taki et al., 2013). Results have been quite variable, sometimes even contradictory (e.g., the volume of the posterior cingulate cortex has been found to be both positively and negatively correlated with agreeableness (Coutinho et al., 2013; DeYoung et al., 2010)). Methodologically, this is in part due to the rather small sample sizes (typically less than 100; 116 in (DeYoung et al., 2010), 52 in (Coutinho et al., 2013)) which undermine replicability (Button et al., 2013); studies with larger sample sizes (W.-Y. Liu et al., 2013) typically fail to replicate previous results.

More recently, surface-based morphometry (SBM) has emerged as a promising measure to study structural brain correlates of personality (Bjørnebekk et al., 2013; Holmes et al., 2012; Rauch et al., 2005; Riccelli, Toschi, Nigro, Terracciano, & Passamonti, 2017; Wright et al., 2006). It has the advantage of disentangling several geometric aspects of brain structure which may contribute to differences detected in VBM, such as cortical thickness (Hutton, Draganski, Ashburner, & Weiskopf, 2009), cortical volume, and folding. Although many studies using SBM are once again limited by small sample sizes, one recent study (Riccelli et al., 2017) used 507 subjects to investigate personality, although it had other limitations (e.g., using a correlational, rather than a predictive framework (Dubois & Adolphs, 2016; Woo, Chang, Lindquist, & Wager, 2017; Yarkoni & Westfall, 2017)).

There is much room for improvement in structural MRI studies of personality traits. The limitation of small sample sizes can now be overcome, since all MRI studies regularly collect structural scans, and recent consortia and data sharing efforts have lead to the accumulation of large publicly available datasets (Job et al., 2017; Miller et al., 2016; Van Essen et al., 2013). One could imagine a mechanism by which personality assessments, if not available already within these datasets, are collected later (Mar, Spreng, & Deyoung, 2013), yielding large samples for relating structural MRI to personality. Lack of out-of-sample generalizability, a limitation of almost all studies that we raised above, can be overcome using cross-validation techniques, or by setting aside a replication sample. In short: despite a considerable historical literature that has investigated the association between personality traits and structural MRI measures, there are as yet no very compelling findings because prior studies have been unable to surmount this list of limitations.

#### Diffusion MRI studies

Several studies have looked for a relationship between white-matter integrity as assessed by DTI and personality factors (Cohen, Schoene-Bake, Elger, & Weber, 2008; Kim & Whalen, 2009; Westlye, Bjørnebekk, Grydeland, Fjell, & Walhovd, 2011; Xu & Potenza, 2012). As with structural MRI studies, extant focal findings often fail to replicate with larger samples of subjects, which tend to find more widespread differences linked to personality traits (Bjørnebekk et al., 2013). The same concerns mentioned in the previous section, in particular the lack of a predictive framework (e.g., using cross-validation), plague this literature; similar recommendations can be made to increase the reproducibility of this line of research, in particular aggregating data (Miller et al., 2016; Van Essen et al., 2013) and using out-of-sample prediction (Yarkoni & Westfall, 2017).

#### Functional MRI studies

Functional MRI (fMRI) measures local changes in blood flow and blood oxygenation as a surrogate of the metabolic demands due to neuronal activity (Logothetis & Wandell, 2004).There are two main paradigms that have been used to relate fMRI data to personality traits: task-based fMRI and resting-state fMRI.

Task-based fMRI studies are based on the assumption that differences in personality may affect information-processing in specific tasks (Yarkoni., 2015). Personality variables are hypothesized to influence cognitive mechanisms, whose neural correlates can be studied with fMRI. For example, differences in neuroticism may materialize as differences in emotional reactivity, which can then be mapped onto the brain (T. Canli et al., 2001). There is a very large literature on task-fMRI substrates of personality, which is beyond the scope of this overview. In general, some of the same concerns we raised above also apply to task-fMRI studies, which typically have even smaller sample sizes (Yarkoni, 2009), greatly limiting power to detect individual differences (in personality or any other behavioral measures). Several additional concerns on the validity of fMRI-based individual differences research apply (Dubois & Adolphs, 2016) and a new challenge arises as well: whether the task used has construct validity for a personality trait.

The other paradigm, resting-state fMRI, offers a solution to the sample size problem, as resting-state data is often collected alongside other data, and can easily be aggregated in large online databases (Biswal et al., 2010; Eickhoff, Nichols, Van Horn, & Turner, 2016; Poldrack & Gorgolewski, 2015; Van Horn & Gazzaniga, 2013). It is the type of data we used in the present paper. Resting-state data does not explicitly engage cognitive processes that are thought to be related to personality traits. Instead, it is used to study correlated self-generated activity between brain areas while a subject is at rest. These correlations, which can be highly reliable given enough data (Finn et al., 2015; Laumann et al., 2015; Noble et al., 2017), are thought to reflect stable aspects of brain organization (Xilin Shen et al., 2017; S. M. Smith et al., 2013). There is a large ongoing effort to link individual variations in functional connectivity (FC) assessed with resting-state fMRI to individual traits and psychiatric diagnosis (for reviews see (Dubois & Adolphs, 2016; Orrù, Pettersson-Yeo, Marquand, Sartori, & Mechelli, 2012; S. M. Smith et al., 2013; Woo et al., 2017).

A number of recent studies have investigated functional connectivity markers from resting-state fMRI and their association with personality traits (Adelstein et al., 2011; Aghajani et al., 2014; Baeken et al., 2014; Beaty et al., 2014, 2016; Gao et al., 2013; Jiao et al., 2017; Lei, Zhao, & Chen, 2013; Pang et al., 2016; Ryan, Sheu, & Gianaros, 2011; Takeuchi et al., 2012; Wu, Li, Yuan, & Tian, 2016). Somewhat surprisingly, these resting-state fMRI studies typically also suffer from low sample sizes (typically less than 100 subjects, usually about 40), and the lack of a predictive framework to assess effect size out-of-sample. One of the best extant datasets, the Human Connectome Project (HCP) has only in the past year reached its full sample of over 1000 subjects, now making large sample sizes readily available. To date, only the exploratory “MegaTrawl” (S. Smith et al., 2016) has investigated personality in this database; we believe that ours is the first comprehensive study of personality on the full HCP dataset, offering very substantial improvements over all prior work.

### Measuring Personality

Although there are a number of different schemes and theories for quantifying personality traits, by far the most common and well validated one, and also the only one available for the Human Connectome Project dataset, is the five-factor solution of personality (aka “The Big Five”). This was originally identified through systematic examination of the adjectives in English language that are used to describe human traits. Based on the hypothesis that all important aspects of human personality are reflected in language, Raymond Cattell applied factor analysis to peer ratings of personality and identified 16 common personality factors (Cattell, 1945). Over the next 3 decades, multiple attempts to replicate Cattell’s study using a variety of methods (e.g. self-description and description of others with adjective lists and behavioral descriptions) agreed that the taxonomy of personality could be robustly described through a five-factor solution (Borgatta, 1964; Fiske, 1949; Norman, 1963; G. M. Smith, 1967; Tupes & Christal, 1961). Since the 1980s, the Big Five has emerged as the leading psychometric model in the field of personality psychology (Goldberg, 1981; Robert R. McCrae & John, 1992). The five factors are commonly termed “openness,” “conscientiousness,” “extraversion,” “agreeableness,” and “neuroticism.”

While the Big Five personality dimensions are not based on an independent theory of personality, and in particular have no basis in neuroscience theories of personality, proponents of the Big Five maintain that they provide the best empirically-based integration of the dominant theories of personality, encompassing the alternative theories of Cattell, Guilford and Eysenck (Amelang & Borkenau, 1982). Self-report questionnaires, such as the NEO-FFI (Robert R. McCrae & Costa, 2004), can be used to reliably assess an individual with respect to these five factors. Even though there remain critiques of the Big Five (Block, 1995; Uher, 2015), its proponents argue that its five factors “are both necessary and reasonably sufficient for describing at a global level the major features of personality” (R. R. McCrae, Costa - American Psychologist, & 1986, 1986).

### The present study

As we emphasized above, personality neuroscience based on MRI data confronts two major challenges. First, nearly all studies to date have been severely underpowered due to small sample sizes (Button et al., 2013; Schönbrodt & Perugini, 2013; Yarkoni, 2009). Second, most studies have failed to use a predictive or replication framework (but see (Deris, Montag, Reuter, Weber, & Markett, 2017)), making their generalizability unclear -- a well-recognized problem in neuroscience studies of individual differences (Dubois & Adolphs, 2016; Gabrieli, Ghosh, & Whitfield-Gabrieli, 2015; Yarkoni & Westfall, 2017). The present paper takes these two challenges seriously by applying a predictive framework, together with a built-in replication, to a large, homogeneous resting-state fMRI dataset. We chose to focus on resting-state fMRI data to predict personality, because this is a predictor that could have better mechanistic interpretation than structural MRI measures (since ultimately it is brain function, not structure, that generates the behavior on the basis of which we can infer personality).

Our dataset, the Human Connectome Project resting-state fMRI data (HCP rs-fMRI) makes available over 1000 well assessed healthy adults. With respect to our study, it provided three types of relevant data: (1) substantial high-quality resting-state fMRI (2 sessions per subject on separate days, each consisting of two 15 minute runs, for 1 hour total); (2) personality assessment for each subject (using the NEO-FFI 2); (3) additional basic cognitive assessment (including fluid intelligence and others), as well as demographic information, which can be assessed as potential confounds.

Our primary question was straightforward: given the challenges noted above, is it possible to find evidence that any personality trait can be reliably predicted from fMRI data, using the best available resting-state fMRI dataset together with the best generally used current analysis methods? If the answer to this question is negative, this might suggest that studies to date that have claimed to find associations between resting-state fMRI and personality are false positives (but of course it would still leave open future positive findings, if more sensitive measures are available). If the answer is positive, it would provide an estimate of the effect size that can be expected in future studies; it would provide initial recommendations for data preprocessing, modeling, and statistical treatment; and it would provide a basis for hypothesis-driven investigations that could focus on particular traits and brain regions. As a secondary aim, we wanted to explore the sensitivity of the results to the details of the analysis used and gain some reassurance that any positive findings would be relatively robust with respect to the details of the analysis; we therefore used a few (well established) combinations of inter-subject alignment, preprocessing, and learning models. This is not intended as a systematic, exhaustive foray into all choices that could be made; such an investigation would be extremely valuable, yet is outside the scope of this work.

## Methods

### Dataset

We used data from a public repository, the 1200 subjects release of the Human Connectome Project (HCP) (Van Essen et al., 2013). The HCP provides MRI data and extensive behavioral assessment from almost 1200 subjects. Acquisition parameters and “minimal” preprocessing of the resting-state fMRI data is described in the original publication (Glasser et al., 2013). Briefly, each subject underwent two sessions of resting-state fMRI on separate days, each session with two separate 15 minute acquisitions generating 1200 volumes (customized Siemens Skyra 3 Tesla MRI scanner, TR = 720 ms, TE = 33 ms, flip angle= 52^°^, voxel size = 2 mm isotropic, 72 slices, matrix = 104 × 90, FOV = 208 mm × 180 mm, multiband acceleration factor = 8). The two runs acquired on the same day differed in the phase encoding direction, left-right and right-left (which leads to differential signal intensity especially in ventral temporal and frontal structures). The HCP data was downloaded in its minimally preprocessed form, i.e. after motion correction, B_0_ distortion correction, coregistration to T_1_-weighted images and normalization to MNI space (the T1w image is registered to MNI space with a FLIRT 12 DOF affine and then a FNIRT nonlinear registration, producing the final nonlinear volume transformation from the subject’s native volume space to MNI space).

### Personality assessment, and personality factors

The 60 item version of the Costa and McCrae Neuroticism/Extraversion/Openness Five Factor Inventory (NEO-FFI), which has shown excellent reliability and validity (Robert R. McCrae & Costa, 2004), was administered to HCP subjects. This measure was collected as part of the Penn Computerized Cognitive Battery (R. C. Gur et al., 2001; Ruben C. Gur et al., 2010). Note that the NEO-FFI was recently updated (NEO-FFI-3, 2010), but the test administered to the HCP subjects is the older version (NEO-FFI-2, 2004).

The NEO-FFI is a self-report questionnaire -- the abbreviated version of the 240-item Neuroticism / Extraversion / Openness Personality Inventory Revised (NEO-PI-R (Paul T. Costa & McCrae, 1992)). For each item, participants reported their level of agreement on a 5-point Likert scale, from strongly disagree to strongly agree.

The Openness, Conscientiousness, Extraversion, Agreeableness and Neuroticism scores are derived by coding each item’s answer (strongly disagree = 0; disagree = 1; neither agree nor disagree = 2; agree = 3; strongly agree = 4) and then reverse coding appropriate items and summing into subscales. As the item scores are available in the database, we recomputed the Big Five scores with the following item coding published in the NEO-FFI 2 manual, where * denotes reverse coding :

- Openness: (3*, 8*, 13, 18*, 23*, 28, 33*, 38*, 43, 48*, 53, 58)
- Conscientiousness: (5, 10, 15*, 20, 25, 30*, 35, 40, 45*, 50, 55*, 60)
- Extraversion: (2, 7, 12*, 17, 22, 27*, 32, 37, 42*, 47, 52, 57*)
- Agreeableness: (4, 9*, 14*, 19, 24*, 29*, 34, 39*, 44*, 49, 54*, 59*)
- Neuroticism: (1*, 6, 11, 16*, 21, 26, 31*, 36, 41, 46*, 51, 56)

We note that the Agreeableness factor score that we calculated was slightly discrepant with the score in the HCP database due to an error in the HCP database in not reverse-coding item 59 at that time (downloaded 06/07/2017). This issue was reported on the HCP listserv (Gray, 2017).

To test the internal consistency of each of the Big Five personality traits in our sample, Cronbach’s alpha was calculated.

Each of the Big Five personality traits can be decomposed into further facets (P. T. Costa Jr & McCrae, 1995), but we did not attempt to predict these facets from our data. Not only does each facet rely on fewer items and thus constitute a noisier measure, which necessarily reduces predictability from neural data (Gignac & Bates, 2017); also, trying to predict many traits leads to a multiple comparison problem which then needs to be accounted for (for an extreme example, see the HCP “MegaTrawl” (S. Smith et al., 2016)).

Despite their theoretical orthogonality, the Big Five are often found to be correlated with one another in typical subject samples. Some authors have suggested that these inter-correlations suggest a higher-order structure, and two superordinate factors have been described in the literature, often referred to as {α / socialization / stability} and {β / personal growth / plasticity} (Blackburn, Renwick, Donnelly, & Logan, 2004; DeYoung, 2006; Digman, 1997). The theoretical basis for the existence of these superordinate factors is highly debated (Robert R. McCrae et al., 2008), and it is not our intention to enter this debate. However, these superordinate factors are less noisy (have lower associated measurement error) than the Big 5, as they are derived from a larger number of test items; this may improve predictability (Gignac & Bates, 2017). Hence, we performed a principal component analysis on the 5 factor scores to extract 2 orthogonal superordinate components, and tested the predictability of these from the HCP rs-fMRI data, in addition to the original five factors.

While we used resting-state fMRI data from two separate sessions (typically collected on consecutive days), there was only a single set of behavioral data available; the NEO-FFI was typically administered on the same day as the second session of resting-state fMRI (Van Essen et al., 2013).

### Fluid intelligence assessment

An estimate of fluid intelligence is available as the *PMAT24_A_CR* measure in the HCP dataset. This proxy for fluid intelligence is based on a short version of Raven’s progressive matrices (24 items) (Bilker et al., 2012); scores are integers indicating number of correct items. We used this fluid intelligence score for two purposes: i) as a benchmark comparison in our predictive analyses, since others have previously reported that this measure of fluid intelligence could be predicted from resting-state fMRI in the HCP dataset (Finn et al., 2015; Noble et al., 2017); ii) as a de-confounding variable (see “Assessment and removal of potential confounds” below). Note that we recently performed a factor analysis of the scores on all cognitive tasks in the HCP to derive a more reliable measure of intelligence; this g-factor could be predicted better than the 24-item score from resting-state data (Dubois, Galdi, Paul, & Adolphs, 2018).

### Subject selection

The total number of subjects in the 1200-subject release of the HCP dataset is N=1206. We applied the following criteria to include/exclude subjects from our analyses (listing in parentheses the HCP database field codes). i) Complete neuropsychological datasets. Subjects must have completed all relevant neuropsychological testing (PMAT_Compl=True, NEO-FFI_Compl=True, Non-TB_Compl=True, VisProc_Compl=True, SCPT_Compl=True, IWRD_Compl=True, VSPLOT_Compl=True) and the Mini Mental Status Exam (MMSE_Compl=True). Any subjects with missing values in any of the tests or test items were discarded. This left us with N = 1183 subjects. ii) Cognitive compromise. We excluded subjects with a score of 26 or below on the MMSE, which could indicate marked cognitive impairment in this highly educated sample of adults under age 40 (Crum, Anthony, Bassett, & Folstein, 1993). This left us with N = 1181 subjects (638 females, 28.8 +/- 3.7 y.o., range 22-37 y.o). Furthermore, iii) subjects must have completed all resting-state fMRI scans (3T_RS-fMRI_PctCompl=100), which leaves us with N = 988 subjects. Finally, iv) we further excluded subjects with a root-mean-squared frame-to-frame head motion estimate (Movement_Relative_RMS.txt) exceeding 0.15mm in any of the 4 resting-state runs (threshold similar to (Finn et al., 2015)). This left us with the final sample of N = 884 subjects (Table S1; 475 females, 28.6 +/- 3.7 y.o., range 22-36 y.o.) for predictive analyses based on resting-state data.

### Assessment and removal of potential confounds

We computed the correlation of each of the personality factors with gender (*Gender*), age (*Age_in_Yrs*, restricted), handedness (*Handedness*, restricted) and fluid intelligence (*PMAT24_A_CR*). We also looked for differences in personality in our subject sample with other variables that are likely to affect FC matrices, such as brain size (we used *FS_BrainSeg_Vol*), motion (we computed the sum of framewise displacement in each run), and the multiband reconstruction algorithm which changed in the third quarter of HCP data collection (*fMRI_3T_ReconVrs*). Correlations are shown in **Figure 2a** below. We then used multiple linear regression to regress these variables from each of the personality scores and remove their confounding effects.

Note that we do not control for differences in cortical thickness and other morphometric features, which have been reported to be correlated with personality factors (e.g. (Riccelli et al., 2017)). These likely interact with FC measures and should eventually be accounted for in a full model, yet this was deemed outside the scope of the present study.

The five personality factors are intercorrelated to some degree (see Results, **Figure 2a**). We did not orthogonalize them-- consequently predictability would be expected also to correlate slightly among personality factors.

It could be argued that controlling for variables such as gender and fluid intelligence risks producing a conservative, but perhaps overly pessimistic picture. Indeed, there are well established gender differences in personality (Feingold, 1994; Schmitt, Realo, Voracek, & Allik, 2008), which might well be based on gender differences in functional connectivity (similar arguments can be made with respect to age (Allemand, Zimprich, & Hendriks, 2008; Soto, John, Gosling, & Potter, 2011) and fluid intelligence (Chamorro-Premuzic & Furnham, 2004; Rammstedt, Danner, & Martin, 2016)). Since the causal primacy of these variables with respect to personality is unknown, it is possible that regressing out sex and age could regress out substantial meaningful information about personality. We therefore also report supplemental results with a less conservative de-confounding procedure -- only regressing out obvious confounds which are not plausibly related to personality, but which would plausibly influence functional connectivity data: image reconstruction algorithm, framewise displacement, and brain size measures.

### Data preprocessing

Resting-state data must be preprocessed beyond “minimal preprocessing”, due to the presence of multiple noise components, such as subject motion and physiological fluctuations. Several approaches have been proposed to remove these noise components and clean the data, however the community has not yet reached a consensus on the “best” denoising pipeline for resting-state fMRI data (Caballero-Gaudes & Reynolds, 2016; Ciric et al., 2017; Murphy & Fox, 2017; Siegel et al., 2016). Most of the steps taken to denoise resting-state data have limitations, and it is unlikely that there is a set of denoising steps that can completely remove noise without also discarding some of the signal of interest. Categories of denoising operations that have been proposed comprise tissue regression, motion regression, noise component regression, temporal filtering and volume censoring. Each of these categories may be implemented in several ways. There exist several excellent reviews of the pros and cons of various denoising steps (Caballero-Gaudes & Reynolds, 2016; T. T. Liu, 2016; Murphy, Birn, & Bandettini, 2013; J. D. Power et al., 2014).

Here, instead of picking a single denoising strategy combining steps used in the previous literature, we set out to explore three reasonable alternatives, which we refer to as A, B, and C (**Figure 1c**). To easily apply these preprocessing strategies in a single framework, using input data that is either volumetric or surface-based, we developed an in-house, Python (v2.7.14)-based pipeline (mostly based on open source libraries and frameworks for scientific computing, including SciPy (v0.19.0), Numpy (v1.11.3), NiLearn (v0.2.6), NiBabel (v2.1.0), Scikit-learn (v0.18.1) (Abraham et al., 2014; K. Gorgolewski et al., 2011; K. J. Gorgolewski et al., 2017; Pedregosa et al., 2011; Walt, Colbert, & Varoquaux, 2011)) implementing the most common denoising steps described in previous literature.

**Figure 1.**
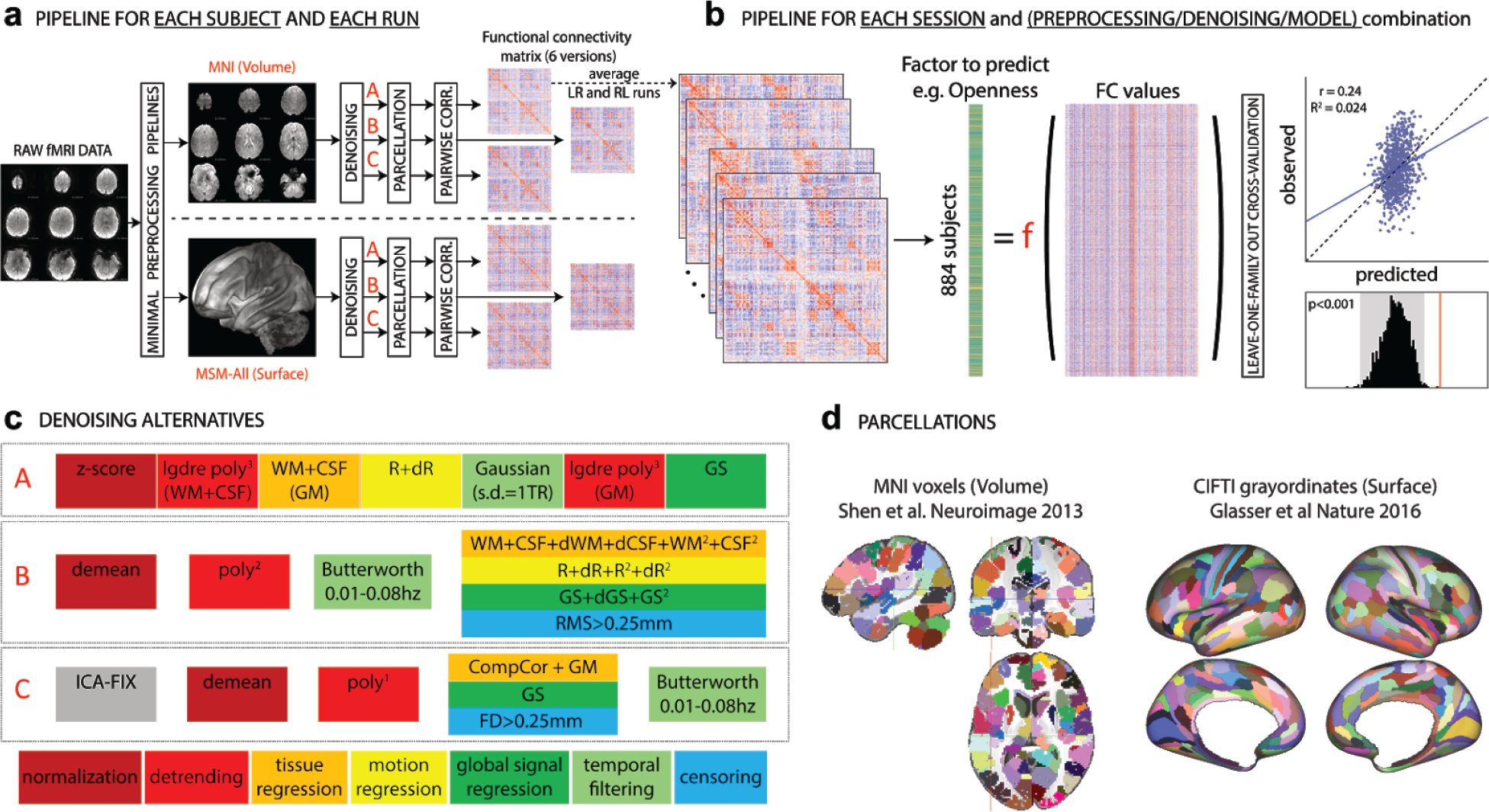
Overview of our approach. In total, we separately analyzed 36 different sets of results: 2 data sessions × 2 alignment/brain parcellation schemes × 3 preprocessing pipelines × 3 predictive models (univariate positive, univariate negative, and multivariate). **a**. The data from each selected HCP subject (N_subjects_=884) and each run (REST1_LR, REST1_RL, REST2_LR, REST2_RL) was downloaded after minimal preprocessing, both in MNI space, and in MSM-All space. The _LR and _RL runs within each session were averaged, producing 2 datasets that we call REST1 and REST2 henceforth. Data for REST1 and REST2, and for both spaces (MNI, MSM-All) were analyzed separately. We applied three alternate denoising pipelines to remove typical confounds found in resting-state fMRI data (see **c**). We then parcellated the data (see **d**) and built a functional connectivity matrix separately for each alternative. This yielded 6 FC matrices per run and per subject. In red: alternatives taken and presented in this paper. **b**. For each of the 6 alternatives, an average FC matrix was computed for REST1 (from REST1_LR and REST1_RL), for REST2 (from REST2_LR and REST2_RL), and for all runs together, REST12. For a given session, we built a (N_subjects_ × N_edges_) matrix, stacking the upper triangular part of all subjects’ FC matrices (the lower triangular part is discarded, because FC matrices are diagonally symmetric). Each column thus corresponds to a single entry in the upper triangle of the FC matrix (a pairwise correlation between two brain parcels, or edge) across all 884 subjects. There are a total of N_parcels_(N_parcels_-1)/2 edges (thus: 35778 edges for the MNI parcellation, 64620 edges for the MSM-All parcellation). This was the data from which we then predicted individual differences in each of the personality factors. We used two different linear models (see text), and a leave-one-family-out cross validation scheme. The final result is a predicted score for each subject, against which we correlate the observed score for statistical assessment of the prediction. Permutations are used to assess statistical significance. **c**. Detail of the three denoising alternatives. These are common denoising strategies for resting-state fMRI. The steps are color-coded to indicate the category of operation they correspond to (legend at the bottom). See text for details. **d**. The parcellations used for the MNI-space and MSM-All space, respectively. Parcels are randomly colored for visualization. Note that the parcellation used for MSM-All space does not include subcortical structures, while the parcellation used for MNI space does.

**Pipeline A** reproduces as closely as possible the strategy described in (Finn et al., 2015) and consists of seven consecutive steps: 1) the signal at each voxel is z-score normalized; 2) using tissue masks, temporal drifts from cerebrospinal fluid (CSF) and white matter (WM) are removed with third-degree Legendre polynomial regressors; 3) the mean signals of CSF and WM are computed and regressed from gray matter voxels; 4) translational and rotational realignment parameters and their temporal derivatives are used as explanatory variables in motion regression; 5) signals are low-pass filtered with a Gaussian kernel with a standard deviation of 1 TR, i.e. 720ms in the HCP dataset; 6) the temporal drift from gray matter signal is removed using a third-degree Legendre polynomial regressor; 7) global signal regression is performed.

**Pipeline B**, described in (Ciric et al., 2017; Satterthwaite, Wolf, et al., 2013), is composed of four steps in our implementation: 1) normalization at voxel-level is performed by subtracting the mean from each voxel’s time series; 2) linear and quadratic trends are removed with polynomial regressors; 3) temporal filtering is performed with a first order Butterworth filter with a passband between 0.01 and 0.08 Hz (after linearly interpolating volumes to be censored, cf. step 4); 4) tissue regression (CSF and WM signals with their derivatives and quadratic terms), motion regression (realignment parameters with their derivatives, quadratic terms and square of derivatives), global signal regression (whole brain signal with derivative and quadratic term), and censoring of volumes with a root-mean-squared (RMS) displacement that exceeded 0.25 mm are combined in a single regression model.

**Pipeline C**, inspired by (Siegel et al., 2016), is implemented as follows: 1) an automated independent component (IC)-based denoising was performed with ICA-FIX (Salimi-Khorshidi et al., 2014). Instead of running ICA-FIX ourselves, we downloaded the FIX-denoised data which is available from the HCP database; 2) voxel signals were demeaned and 3) detrended with a first degree polynomial; 4) CompCor, a PCA-based method proposed by (Behzadi, Restom, Liau, & Liu, 2007) was applied to derive 5 components from CSF and WM signals; these were regressed out of the data, together with gray matter and whole-brain mean signals; volumes with a framewise displacement greater than 0.25 mm or a variance of differentiated signal (DVARS) greater than 105% of the run median DVARS were discarded as well; 5) temporal filtering was performed with a first-order Butterworth band-pass filter between 0.01 and 0.08 Hz, after linearly interpolating censored volumes.

### Inter-subject alignment, parcellation, and functional connectivity matrix generation

An important choice in processing fMRI data is how to align subjects in the first place. The most common approach is to warp individual brains to a common volumetric template, typically MNI152. However, cortex is a 2D structure; hence, surface-based algorithms that rely on cortical folding to map individual brains to a template may be a better approach. Yet another improvement in aligning subjects may come from using functional information alongside anatomical information - this is what the multimodal surface matching (MSM) framework achieves (Robinson et al., 2014). MSM-All aligned data, in which intersubject registration uses individual cortical folding, myelin maps, and resting-state fMRI correlation data, is available for download from the HCP database.

Our prediction analyses below are based on functional connectivity matrices. While voxel- (or vertex-) wise functional connectivity matrices can be derived, their dimensionality is too high compared to the number of examples in the context of a machine-learning based predictive approach. PCA or other dimensionality reduction applied to the voxelwise data can be used, but this often comes at the cost of losing neuroanatomical specificity. Hence, we work with the most common type of data: parcellated data, in which data from many voxels (or vertices) is aggregated anatomically and the signal within a parcel is averaged over its constituent voxels. Choosing a parcellation scheme is the first step in a network analysis of the brain (Sporns, 2013), yet once again there is no consensus on the “best” parcellation. There are two main approaches to defining network nodes in the brain: nodes may be a set of overlapping, weighted masks, e.g. obtained using independent component analysis (ICA) of BOLD fMRI data (S. M. Smith et al., 2013); or a set of discrete, non-overlapping binary masks, also known as a hard parcellation (Glasser, Coalson, et al., 2016; Gordon et al., 2014). We chose to work with a hard parcellation, which we find easier to interpret.

Here we present results based on a classical volumetric alignment, together with a volumetric parcellation of the brain into 268 nodes (Finn et al., 2015; X. Shen, Tokoglu, Papademetris, & Constable, 2013); and, for comparison, results based on MSM-All data, together with a parcellation that was specifically derived from this data (Glasser, Coalson, et al., 2016) (**Figure 1d**).

Timeseries extraction simply consisted in averaging data from voxels (or grayordinates) within each parcel, and matrix generation in pairwise correlating parcel time series (Pearson correlation coefficient). FC matrices were averaged across runs (all averaging used Fisher-z transforms) acquired with left-right and right-left phase encoding in each session, i.e. we derived two FC matrices per subject, one for REST1 (from REST1_LR and REST1_RL) and one for REST2 (from REST2_LR and REST2_RL); we also derived a FC matrix averaged across all runs (REST12).

### Test-retest comparisons

We applied all three denoising pipelines to the data of all subjects. We then compared the functional connectivity (FC) matrices produced by each of these strategies, using several metrics. One metric that we used follows from the connectome fingerprinting work of (Finn et al., 2015), and was recently labeled the identification success rate (ISR) (Noble et al., 2017). Identification of subject S is successful if, out of all subjects’ FC matrices derived from REST2, subject S’s is the most highly correlated with subject S’s FC matrix from REST1 (identification can also be performed from REST2 to REST1; results are very similar). The ISR gives an estimate of the reliability and specificity of the entire FC matrix at the individual subject level, and is influenced both by within-subject test-retest reliability as well as by discriminability amongst all subjects in the sample. Relatedly, it is desirable to have similarities (and differences) between all subjects be relatively stable across repeated testing sessions. Following an approach introduced in (Geerligs, Rubinov, Cam-Can, & Henson, 2015), we computed the pairwise similarity between subjects separately for session 1 and session 2, constructing a N_subjects_×N_subjects_ matrix for each session. We then compared these matrices using a simple Pearson correlation. Finally, we used a metric targeted at behavioral utility, and inspired by (Geerligs, Rubinov, et al., 2015): for each edge (the correlation value between a given pair of brain parcels) in the FC matrix, we computed its correlation with a stable trait across subjects, and built a matrix representing the relationship of each edge to this trait, separately for session 1 and session 2. We then compared these matrices using a simple Pearson correlation. The more edges reliably correlate with the stable trait, the higher the correlation between session 1 and session 2 matrices. It should be noted that trait stability is an untested assumption with this approach, because in fact only a single trait score was available in the HCP, collected at the time of session 2. We performed this analysis for the measure of fluid intelligence available in the HCP (*PMAT24_A_CR*) as well as all Big Five personality factors.

### Prediction models

There is no obvious “best” model available to predict individual behavioral measures from functional connectivity data (Abraham et al., 2016). So far, most attempts have relied on linear machine learning approaches. This is partly related to the well-known “curse of dimensionality”: despite the relatively large sample size that is available to us (N=884 subjects), it is still about an order of magnitude smaller than the typical number of features included in the predictive model. In such situations, fitting relatively simple linear models is less prone to overfitting than fitting complex nonlinear models.

There are several choices of linear prediction models. Here, we present the results of two methods that have been used in the literature for similar purposes: **1)** a simple, “univariate” regression model as used in (Finn et al., 2015), and further advocated by (Xilin Shen et al., 2017), preceded by feature selection; and **2)** a regularized linear regression approach, based on elastic-net penalization (Zou & Hastie, 2005). We describe each of these in more detail next.

**Model (1)** is the simplest model, and the one proposed by (Finn et al., 2015), consisting in a univariate regressor where the dependent variable is the score to be predicted and the explanatory variable is a scalar value that summarizes the functional connectivity network strength (i.e., the sum of edge weights). A filtering approach is used to select features (edges in the FC correlation matrix) that are correlated with the behavioral score on the training set: edges that correlate with the behavioral score with a p-value less than 0.01 are kept. Two distinct models are built using edges of the network that are positively and negatively correlated with the score, respectively. This method has the advantage of being extremely fast to compute, but some main limitations are that i) it condenses all the information contained in the connectivity network into a single measure and does not account for any interactions between edges and ii) it arbitrarily builds two separate models (one for positively correlated edges, one for negatively correlated edges; they are referred to as the positive and the negative models (Finn et al., 2015)) and does not offer a way to integrate them. We report results from both the positive and negative models for completeness.

To address the limitations of the univariate model(s), we also included a multivariate model. **Model (2)** kept the same filtering approach as for the univariate model (discard edges for which the p-value of the correlation with the behavioral score is greater than 0.01); this choice allows for a better comparison of the multivariate and univariate models, and for faster computation. Elastic Net is a regularized regression method that linearly combines L1- (lasso) and L2- (ridge) penalties to shrink some of the regressor coefficients toward zero, thus retaining just a subset of features. The lasso model performs continuous shrinkage and automatic variable selection simultaneously, but in the presence of a group of highly correlated features, it tends to arbitrarily select one feature from the group. With high-dimensional data and few examples, the ridge model has been shown to outperform lasso; yet it cannot produce a sparse model since all the predictors are retained. Combining the two approaches, elastic net is able to do variable selection and coefficient shrinkage while retaining groups of correlated variables. Here however, based on preliminary experiments and on the fact that it is unlikely that just a few edges contribute to prediction, we fixed the L1 ratio (which weights the L1- and L2- regularizations) to 0.01, which amounts to almost pure ridge regression. We used 3-fold nested cross-validation (with balanced “classes”, based on a partitioning of the training data into quartiles) to choose the alpha parameter (among 50 possible values) that weighs the penalty term.

### Cross-validation scheme

In the HCP dataset, several subjects are genetically related (in our final subject sample, there were 410 unique families). To avoid biasing the results due to this family structure (e.g., perhaps having a sibling in the training set would facilitate prediction for a test subject), we implemented a leave-one-family-out cross-validation scheme for all predictive analyses.

### Statistical assessment of predictions

Several measures can be used to assess the quality of prediction. A typical approach is to plot observed vs. predicted values (rather than predicted vs. observed (Piñeiro, Perelman, Guerschman, & Paruelo, 2008)). The Pearson correlation coefficient between observed scores and predicted scores is often reported as a measure of prediction (e.g. (Finn et al., 2015)), given its clear graphical interpretation. However, in the context of cross-validation, it is incorrect to square this correlation coefficient to obtain the coefficient of determination R^2^, which is often taken to reflect the proportion of variance explained by the model (Alexander, Tropsha, & Winkler, 2015); instead, the coefficient of determination R^2^ should be calculated as:

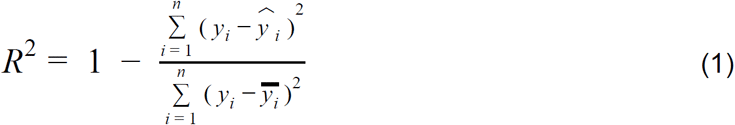

where *n* is the number of observations (subjects), y is the observed response variable, 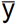 is its mean, and 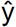 is the corresponding predicted value. Equation (1) therefore measures the size of the residuals from the model compared with the size of the residuals for a null model where all of the predictions are the same, i.e., the mean value 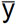. In a cross-validated prediction context, R^2^ can actually take negative values (in cases when the denominator is larger than the numerator, i.e. when the sum of squared errors is larger than that of the null model)! Yet another, related statistic to evaluate prediction outcome is the Root Mean Square Deviation (RMSD), defined in (Piñeiro et al., 2008) as:

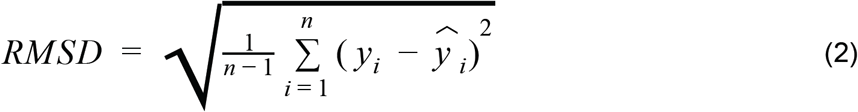

RMSD as defined in (2) represents the standard deviation of the residuals. To facilitate interpretation, it can be normalized by dividing it by the standard deviation of the observed values:

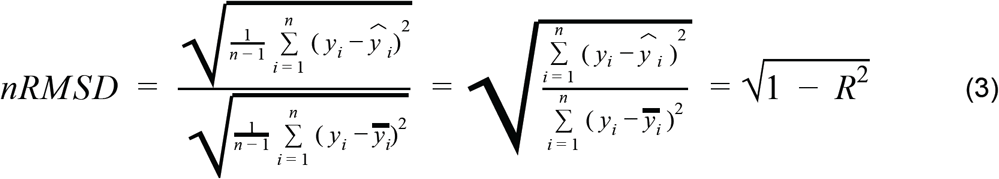

nRMSD thus has a very direct link to R^2^ (3); it is interpretable as the average deviation of each predicted value to the corresponding observed value, and is expressed as a fraction of the standard deviation of the observed values.

In a cross-validation scheme, the folds are not independent of each other. This means that statistical assessment of the cross-validated performance using parametric statistical tests is problematic (Combrisson & Jerbi, 2015; Noirhomme et al., 2014). Proper statistical assessment should thus be done using permutation testing on the actual data. To establish the empirical distribution of chance, we ran our final predictive analyses using 1000 random permutations of the scores (shuffling scores randomly between subjects, keeping everything else exactly the same, including the family structure).

## Results

### Characterization of behavioral measures

#### Internal consistency, distribution, and inter-correlations of personality traits

In our final subject sample (N=884), there was good internal consistency for each personality trait, as measured with Cronbach’s α. We found: Openness, α = 0.76; Conscientiousness α = 0.81; Extraversion, α = 0.78; Agreeableness, α = 0.76; and Neuroticism, α = 0.85. These compare well with the values reported by (Robert R. McCrae & Costa, 2004).

Scores on all factors were nearly normally distributed by visual inspection, although the null hypothesis of a normal distribution was rejected for all but Agreeableness (using D’Agostino and Pearson’s (D’Agostino & Pearson, 1973) normality test as implemented in SciPy) (**Figure 2b**).

**Figure 2.**
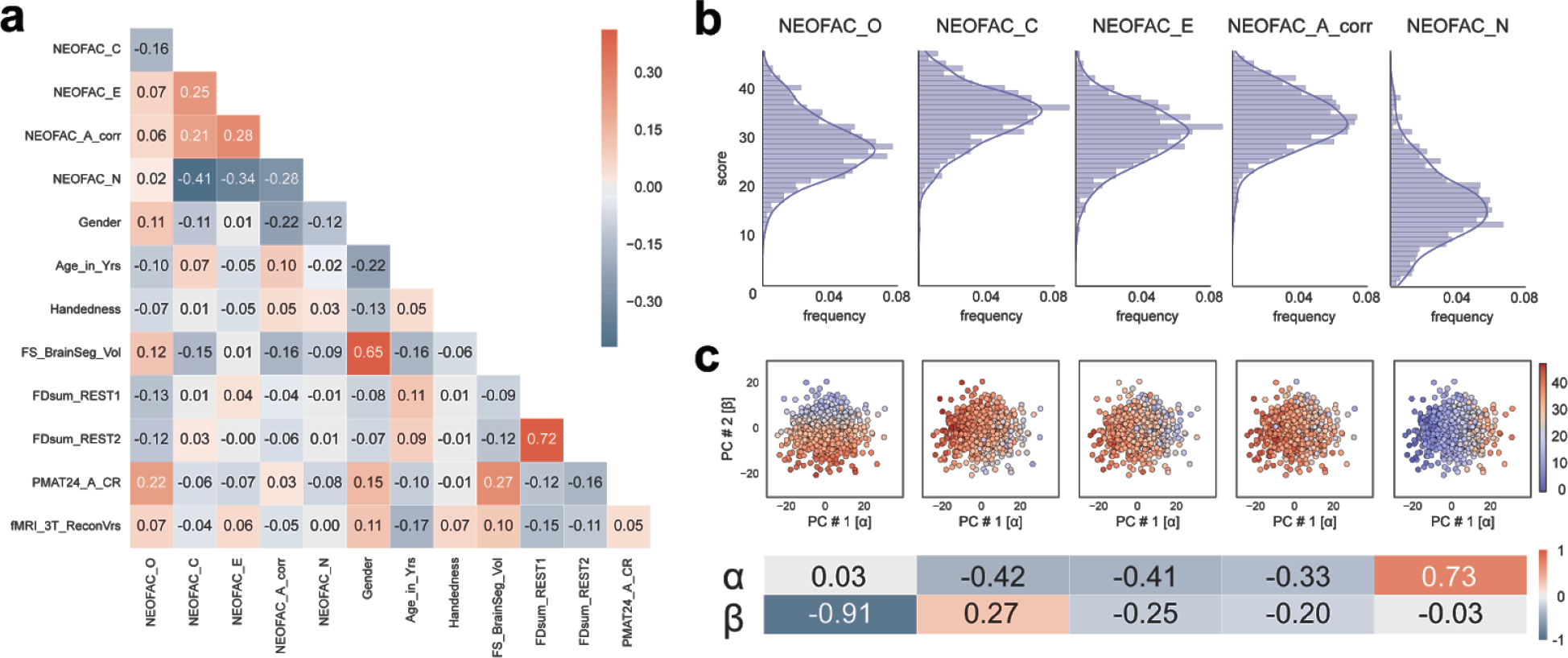
Structure of Personality factors in our subject sample (N=884). **a.** The five personality factors were not orthogonal in our sample. Neuroticism was anticorrelated with Conscientiousness, Extraversion and Agreeableness, and the latter three were positively correlated with each other. Openness correlated more weakly with other factors. There were highly significant correlations with other behavioral and demographic variables, which we accounted for in our subsequent analyses by regressing them out of the personality scores (see next section). **b.** Distributions of the five personality scores in our sample. Each of the five personality scores was approximately normally distributed by visual inspection. **c.** Two-dimensional principal component projection; the value for each personality factor in this projection is represented by the color of the dots. The weights for each personality factor are shown at the bottom.

Although in theory the Big Five personality traits should be orthogonal, their estimation from the particular item scoring of versions of the NEO in practice deviates considerably from orthogonality. This intercorrelation amongst the five factors has been reported for the NEO-PI-R (Block, 1995; Saucier, 2002), the NEO-FFI (Block, 1995; Egan, Deary, & Austin, 2000), and alternate instruments (DeYoung, 2006) (but, see (Robert R. McCrae et al., 2008)). Indeed, in our subject sample, we found that the five personality factors were correlated with one another (**Figure 2a**). For example, Neuroticism was anticorrelated with Conscientiousness (r = −0.41, p <10^−37^), Extraversion (r = −0.34, p < 10^−25^), and Agreeableness (r = −0.28, p <10^−16^), while these latter three factors were positively correlated with one another (all r>0.21). Though the theoretical interpretation of these inter-correlations in terms of higher-order factors of personality remains a topic of debate (DeYoung, 2006; Digman, 1997; Robert R. McCrae et al., 2008), we derived two orthogonal higher-order personality dimensions using a principal component analysis of the Big 5 factor scores; we labeled the two derived dimensions α and β, following (Digman, 1997). The first component [α] accounted for 40.3% of the variance, and the second [β] for 21.6% (total variance explained by the two-dimensional PC solution was thus 61.9%). **Figure 2c** shows how the Big Five project on this two-dimensional solution, and the PC loadings.

#### Confounding variables

There are known effects of gender (Ruigrok et al., 2014; Trabzuni et al., 2013), age (Dosenbach et al., 2010; Geerligs, Renken, Saliasi, Maurits, & Lorist, 2015), handedness (Pool, Rehme, Eickhoff, Fink, & Grefkes, 2015), in-scanner motion (J. D. Power, Barnes, Snyder, Schlaggar, & Petersen, 2012; Satterthwaite, Elliott, et al., 2013; Tyszka, Kennedy, Paul, & Adolphs, 2014), brain size (Hänggi, Fövenyi, Liem, Meyer, & Jäncke, 2014) and fluid intelligence (Cole, Yarkoni, Repovs, Anticevic, & Braver, 2012; Finn et al., 2015; Noble et al., 2017) on the functional connectivity patterns measured in the resting-state with fMRI. It is thus necessary to control for these variables: indeed, if a personality factor is correlated with gender, one would be able to predict some of the variance in that personality factor solely from functional connections that are related to gender. The easiest way (though perhaps not the best way, see (Westfall & Yarkoni, 2016)) to control for these confounds is by regressing the confounding variables on the score of interest in our sample of subjects.

We characterized the relationship between each of the personality factors and each of the confounding variables listed above in our subject sample (**Figure 2a**). All personality factors but Extraversion were correlated with gender: women scored higher on Conscientiousness, Agreeableness and Neuroticism, while men scored higher on Openness. In previous literature, women have been reliably found to score higher on Neuroticism and Agreeableness, which we replicated here, while other gender differences are generally inconsistent at the level of the factors (Paul T. Costa, Terracciano, & McCrae, 2001; Feingold, 1994; Weisberg, Deyoung, & Hirsh, 2011). Agreeableness and Openness were significantly correlated with age in our sample, despite our limited age range (22-36 y.o.): younger subjects scored higher on Openness, while older subjects scored higher on Agreeableness. The finding for Openness does not match previous reports (Allemand et al., 2008; Soto et al., 2011), but this may be confounded by other factors such as gender, as our analyses here do not use partial correlations. Motion, quantified as the sum of frame-to-frame displacement over the course of a run (and averaged separately for REST1 and REST2) was correlated with Openness: subjects scoring lower on Openness moved more during the resting-state. Note that motion in REST1 was highly correlated (r=0.72, p<10^−143^) with motion in REST2, indicating that motion itself may be a stable trait, and correlated with other traits. Brain size, obtained from Freesurfer during the minimal preprocessing pipelines, was found to be significantly correlated with all personality factors but Extraversion. Fluid intelligence was positively correlated with Openness, and negatively correlated with Conscientiousness, Extraversion, and Neuroticism, consistent with other reports (Bartels et al., 2012; Chamorro-Premuzic & Furnham, 2004). While the interpretation of these complex relationships would require further work outside the scope of this study, we felt that it was critical to remove shared variance between each personality score and the primary confounding variables before proceeding further. This ensures that our model is trained specifically to predict personality, rather than confounds that covary with personality, although it may also reduce power by removing shared variance (thus providing a conservative result).

Another possible confound, specific to the HCP dataset, is a difference in the image reconstruction algorithm between subjects collected prior to and after April 2013. The reconstruction version leaves a notable signature on the data that can make a large difference in the final analyses produced (Elam, 2015). We found a significant correlation with the Openness factor in our sample. This indicates that the sample of subjects who were scanned with the earlier reconstruction version happened to score slightly less high for the Openness factor than the sample of subjects who were scanned with the later reconstruction version (purely by sampling chance); this of course is meaningless, and a simple consequence of working with finite samples. Therefore, we also included the reconstruction factor as a confound variable.

Importantly, the multiple linear regression used for removing the variance shared with confounds was performed on training data only (in each cross-validation fold during the prediction analysis), and then the fitted weights were applied to both the training and test data. This is critical to avoid any leakage of information, however negligible, from the test data into the training data.

Authors of the HCP-MegaTrawl have used transformed variables (Age^2^) and interaction terms (Gender × Age, Gender × Age^2^) as further confounds (S. Smith et al., 2016). After accounting for the confounds described above, we did not find sizeable correlations with these additional terms (all correlations <0.008), and thus we did not use these additional terms in our confound regression.

### Preprocessing affects test-retest reliability of FC matrices

As we were interested in relatively stable traits (which are unlikely to change much between sessions REST1 and REST2), one clear goal for the denoising steps applied to the minimally preprocessed data was to yield functional connectivity matrices that are as “similar” as possible across the two sessions. We computed several metrics (see Methods) to assess this similarity for each of our three denoising strategies (A, B, and C; cf **Figure 1c**). Of course, no denoising strategy would achieve perfect test-retest reliability of FC matrices since, in addition to inevitable measurement error, the two resting-state sessions for each subject likely feature somewhat different levels of states such as arousal and emotion.

In general, differences in test-retest reliability across metrics were small when comparing the three denoising strategies. Considering the entire FC matrix, the Identification Success Rate (ISR) (Finn et al., 2015; Noble et al., 2017) was high for all strategies, and highest for pipeline B **(Figure 3a)**. The multivariate pairwise distances between subjects were also best reproduced across sessions by pipeline B **(Figure 3b)**. In terms of behavioral utility, i.e. reproducing the pattern of correlations of the different edges with a behavioral score, pipeline A outperformed the others **(Figure 3c)**. All three strategies appear to be reasonable choices, and we would thus expect a similar predictive accuracy under each of them, if there is information about a given score in the functional connectivity matrix.

**Figure 3.**
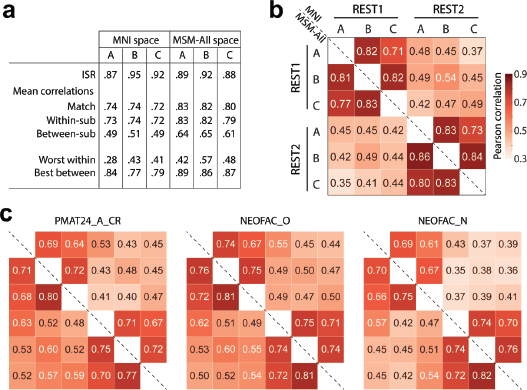
Test-retest comparisons between spaces and denoising strategies. **a**. Identification success rate, and other statistics related to connectome fingerprinting (Finn et al., 2015; Noble et al., 2017). All pipelines had a success rate superior to 87% for identifying the functional connectivity matrix of a subject in REST2 (out of N=884 choices) based on their functional connectivity matrix in REST1. Pipeline B slightly outperformed the others. **b**. Test-retest of the pairwise similarities (based on Pearson correlation) between all subjects (Geerligs, Rubinov, et al., 2015). Overall, for the same session, the three pipelines gave similar pairwise similarities between subjects. About 25% of the variance in pairwise distances was reproduced in REST2, with pipeline B emerging as the winner (0.54^2^=29%). **c**. Test-retest reliability of behavioral utility, quantified as the pattern of correlations between each edge and a behavioral score of interest (Geerligs, Rubinov, et al., 2015). Shown are fluid intelligence, Openness to experience, and Neuroticism (all de-confounded, see main text). Pipeline A gave slightly better test-retest reliability for all behavioral scores. MSM-All outperformed MNI alignment. Neuroticism showed lower test-retest reliability than fluid intelligence or Openness to experience.

We note here already that Neuroticism stands out as having lower test-retest reliability in terms of its relationship to edge values across subjects (**Figure 3c**). This may be a hint that the FC matrices do not carry information about Neuroticism.

### Prediction of fluid intelligence (*PMAT24_A_CR*)

It has been reported that a measure of fluid intelligence, the raw score on a 24-item version of the Raven’s Progressive Matrices (*PMAT24_A_CR*), could be predicted from FC matrices in previous releases of the HCP dataset (Finn et al., 2015; Noble et al., 2017). We generally replicated this result qualitatively for the de-confounded fluid intelligence score (removing variance shared with gender, age, handedness, brain size, motion, and reconstruction version), using a leave-one-family-out cross-validation approach. We found positive correlations across all 36 of our result datasets: 2 sessions × 3 denoising pipelines (A, B & C) × 2 parcellation schemes (in volumetric space and in MSM-All space) × 3 models (univariate positive, univariate negative, and multivariate learning models) (**Figure 4a**, **Table 1**). We note however that, using MNI space and denoising strategy A as in (Finn et al., 2015), the prediction score was very low (REST1: r=0.04; REST2: r=0.03). One difference is that the previous study did not use de-confounding, hence some variance from confounds may have been used in the predictions; also the sample size was much smaller in (Finn et al., 2015) (N=118; but N=606 in (Noble et al., 2017)), and family structure was not accounted for in the cross-validation. We generally found that prediction performance was better in MSM-All space (**Figure 4a**, **Table 1**).

**Table 1.**
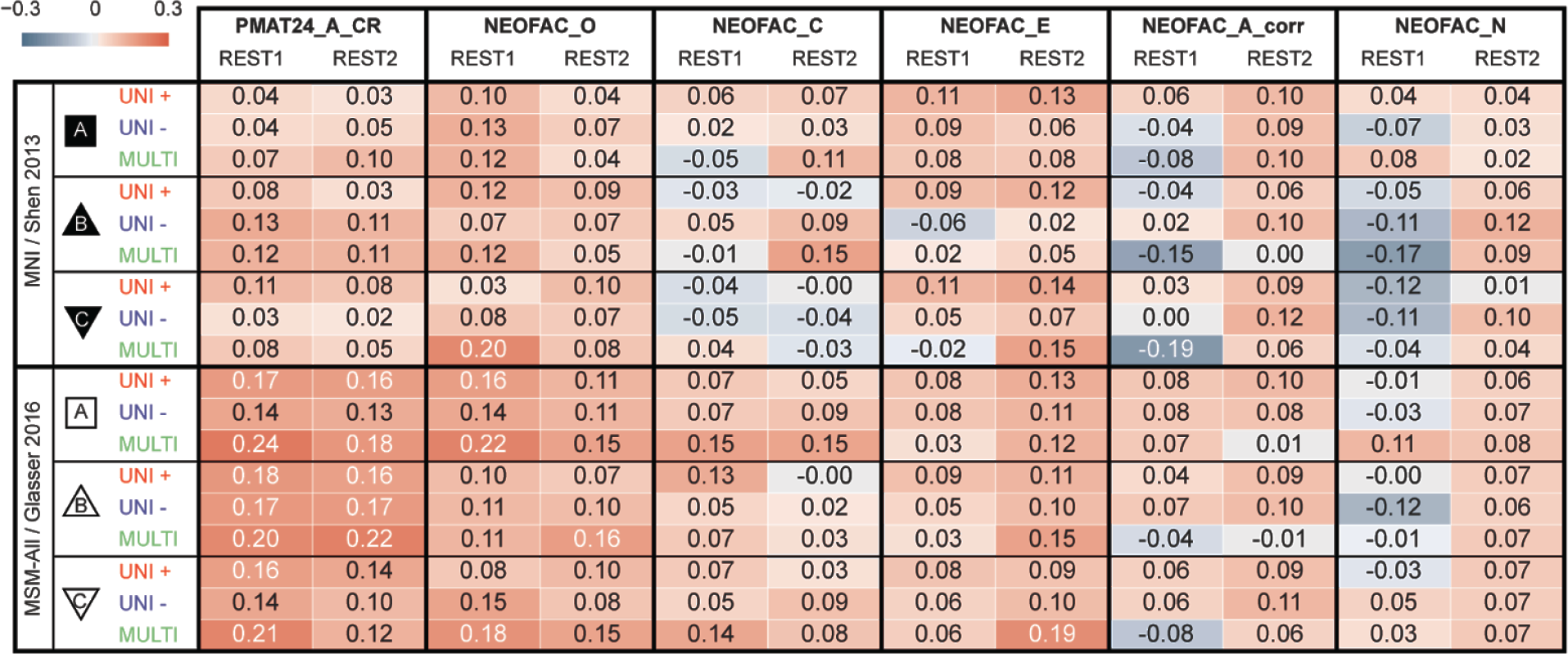
Test-retest prediction results using deconfounded scores. Listed are Pearson correlation coefficients between predicted and observed individual scores, for all behavioral scores and analytical alternatives (the two columns for each score correspond to the two resting-state sessions). See **Supplementary Figure 1** for results with minimal deconfounding.

**Figure 4.**
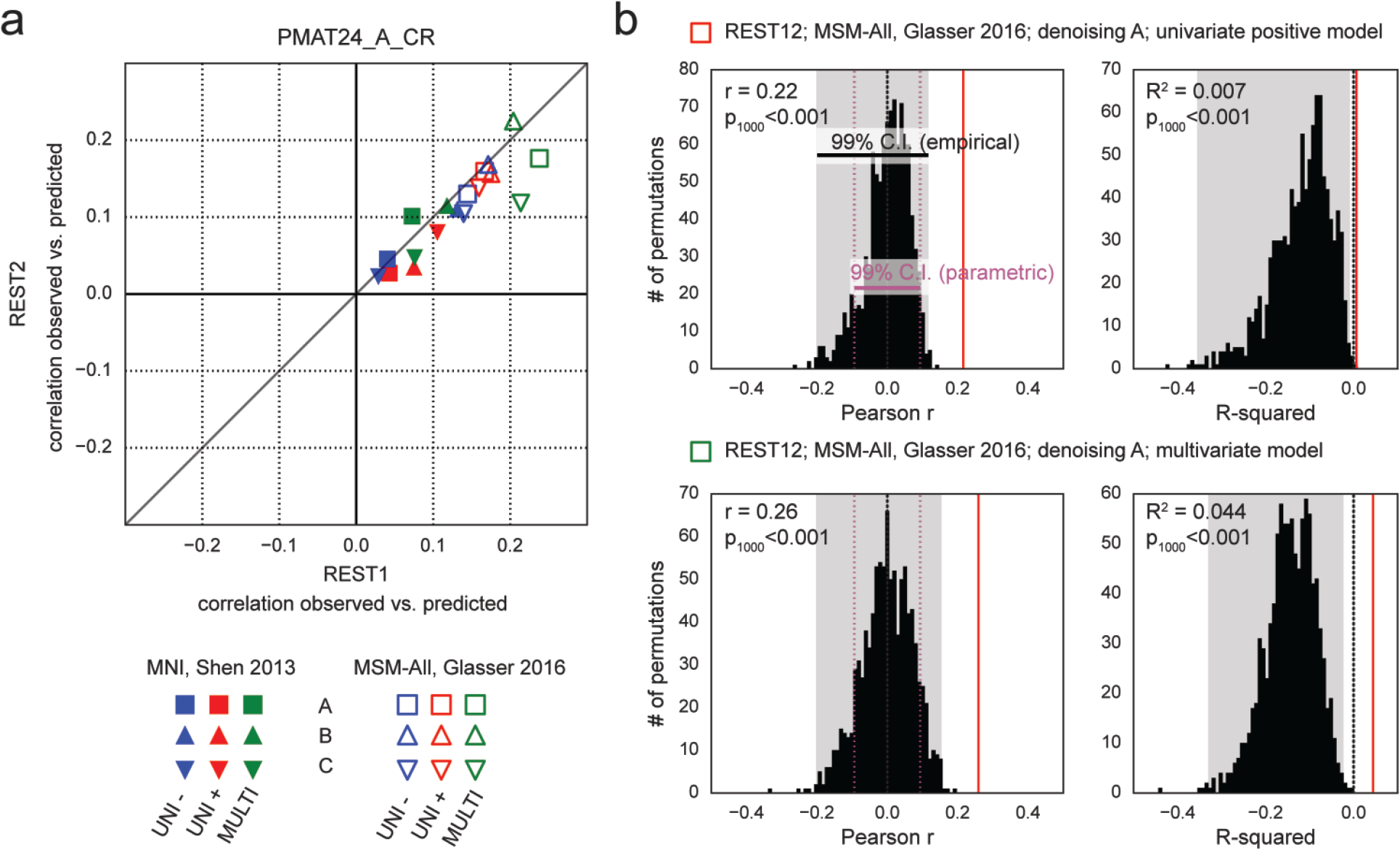
Prediction results for de-confounded fluid intelligence (*PMAT24_A_CR*). **a**. All predictions were assessed using the correlation between the observed scores (the actual scores of the subjects) and the predicted scores. This correlation obtained using the REST2 dataset was plotted against the correlation from the REST1 dataset, to assess test-retest reliability of the prediction outcome. Results in MSM-All space outperformed results in MNI space. The multivariate model slightly outperformed the univariate models (positive and negative). Our results generally showed good test-retest reliability across sessions, although REST1 tended to produce slightly better predictions than REST2. Pearson correlation scores for the predictions are listed in **Table 1**. **Supplementary Figure 1** shows prediction scores with minimal deconfounding. **b**. We ran a final prediction using combined data from all resting-state runs (REST12), in MSM-All space with denoising strategy A (results are shown as vertical red lines). We randomly shuffled the PMAT24_A_CR scores 1000 times while keeping everything else the same, for the univariate model (positive, top) and the multivariate model (bottom). The distribution of prediction scores (Pearson r, and R^2^) under the null hypothesis is shown (black histograms). Note that the empirical 99% confidence interval (shaded gray area) is wider than the parametric CI (shown for reference, magenta dotted lines), and features a heavy tail on the left side for the univariate model. This demonstrates that parametric statistics are not appropriate in the context of cross-validation. Such permutation testing may be computationally prohibitive for more complex models, yet since the chance distribution is model-dependent, it must be performed for statistical assessment.

To generate a final prediction, we combined data from all four resting-state runs (REST12). We chose to use pipeline A and MSM-All space, which we had found to yield the best test-retest reliability in terms of behavioral utility (**Figure 3c**). We obtained r=0.22 (R^2^=0.007, nRMSD=0.997) for the univariate positive model, r=0.18 (R^2^=-0.023, nRMSD=1.012) for the univariate negative model, and r=0.26 (R^2^=0.044, nRMSD=0.978) for the multivariate model. Interestingly, these performances on combined data outperformed performance on REST1 or REST2 alone, suggesting that decreasing noise in the neural data boosts prediction performance. For statistical assessment of predictions, we estimated the distribution of chance for the prediction score under both the univariate positive and the multivariate models, using 1000 random permutations of the subjects’ fluid intelligence scores (**Figure 4b**). For reference we also show parametric statistics thresholds for the correlation coefficients; we found that parametric statistics underestimate the confidence interval for the null hypothesis, hence overestimate significance. Interestingly, the null distributions differed between the univariate and the multivariate models: while the distribution under the multivariate model was roughly symmetric about 0, the distribution under the univariate model was asymmetric with a long tail on the left. The empirical, one-tailed p-values for REST12 MSM-All space data denoised with strategy A and using the univariate positive model, and using the multivariate model, both achieved p<0.001 (none of the 1000 random permutations resulted in a higher prediction score).

### Prediction of the Big Five

We established that our approach reproduces and improves on the previous finding that fluid intelligence can be predicted from resting-state functional connectivity (Finn et al., 2015; Noble et al., 2017). We next turned to predicting each of the Big Five personality factors using the same approach (including de-confounding, which in this case removes variance shared with gender, age, handedness, brain size, motion, reconstruction version and, importantly, fluid intelligence).

Test-retest results across analytical choices are shown in **Figure 5a**, and in **Table 1**. Predictability was lower than for fluid intelligence (*PMAT24_A_CR*) for all Big Five personality factors derived from the NEO-FFI. Openness to experience showed the highest predictability overall, and also the most reproducible across sessions; prediction of Extraversion was moderately reproducible; in contrast, the predictability of the other three personality factors (Agreeableness and Neuroticism, and Conscientiousness) was low and lacked reproducibility.

**Figure 5.**
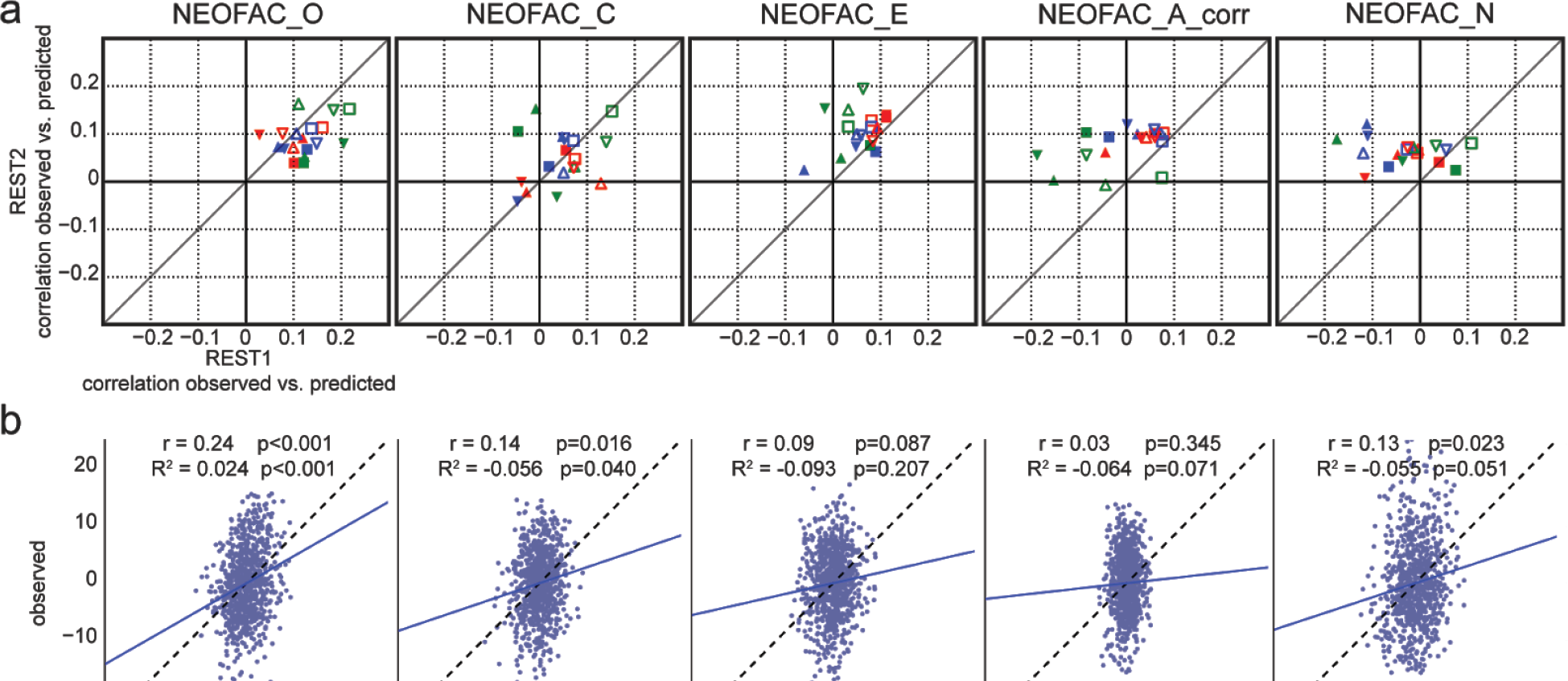
Prediction results for the Big Five personality factors. **a**. Test-retest prediction results for each of the Big Five. Representation is the same as in **Figure 4a**. The only factor that showed consistency across parcellation schemes, denoising strategies, models and sessions was Openness (*NEOFAC_O*), although Extraversion (*NEOFAC_E*) also showed substantial positive correlations. See also **Table 1**. **b**. Prediction results for each of the (demeaned and deconfounded) Big Five, from REST12 FC matrices, using MSM-All inter-subject alignment, denoising strategy A, and the multivariate prediction model. The blue line shows the best fit to the cloud of points (its slope should be close to 1 (dotted line) for good predictions (Piñeiro et al., 2008)). The variance of predicted values is noticeably smaller than the variance of observed values.

**Figure 6.**
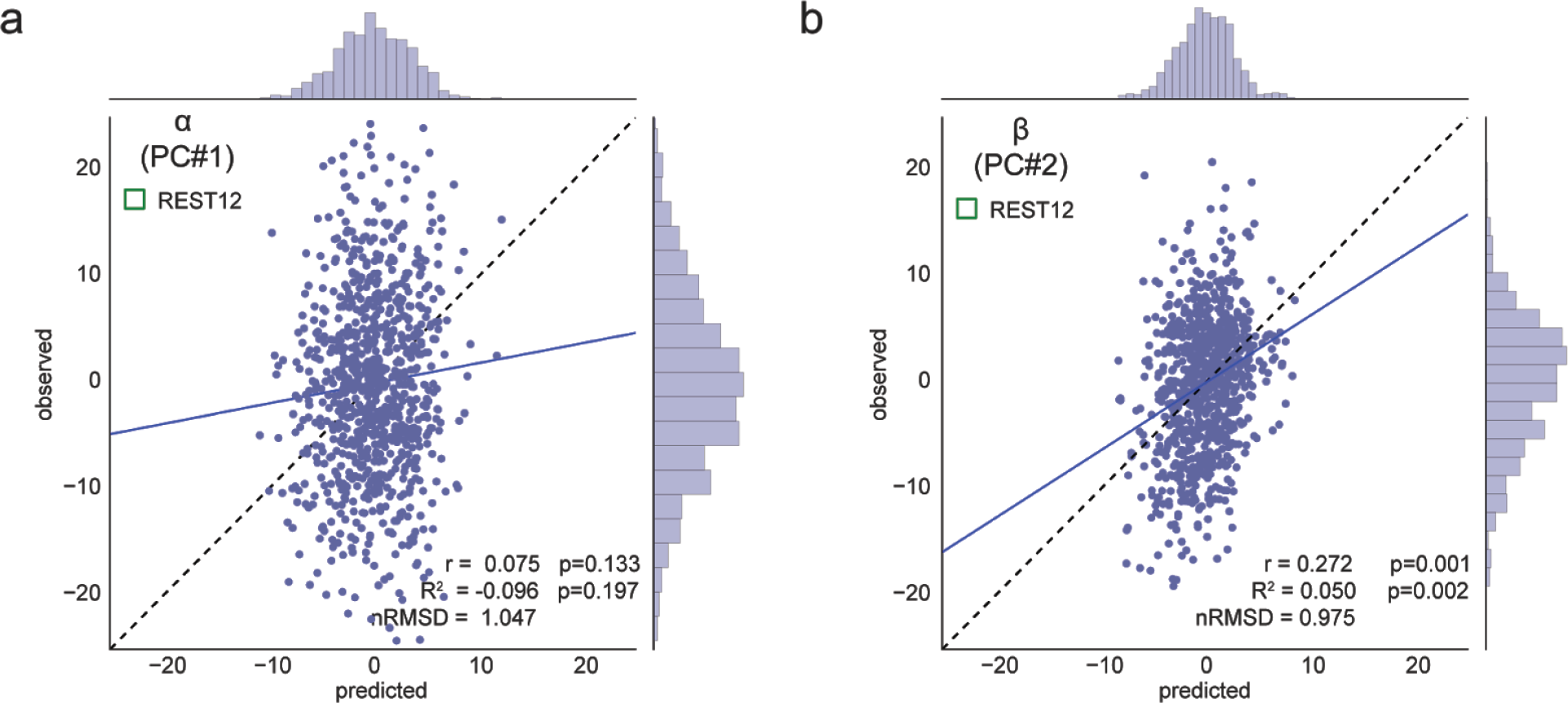
Prediction results for superordinate factors/principal components α and β, using REST12 data (1h of resting-state fMRI per subject). These results use MSM-All inter-subject alignment, denoising strategy A, and the multivariate prediction model. As in **Figure 5b**, the range of predicted scores is much narrower than the range of observed scores. The first PC, α, is not predicted better than chance. α loads mostly on Neuroticism (see **Figure 2c**), which was itself not predicted well (cf. Figure 5). **b.** We can predict about 5% of the variance in the score on the second PC, β. This is better than chance, as established by permutation statistics (p<0.002). β loads mostly on Openness to Experience (see **Figure 2c**), which showed good predictability in the previous section.

It is worth noting that the NEO-FFI test was administered closer in time to REST2 than to REST1 on average; hence one might expect REST2 to yield slightly better results, if the NEO-FFI factor scores reflect a state component. We found that REST2 produced better predictability than REST1 for Extraversion (results fall mostly to the left of the diagonal line of reproducibility), while REST1 produced better results for Openness, hence the data does not reflect our expectation of state effects on predictability.

Although we conducted 18 different analyses for each session with the intent to present all of them in a fairly unbiased manner, it is notable that certain combinations produced the best predictions across different personality scores - some of the same combinations that yielded the best predictability for fluid intelligence (**Figure 4**, above). While the findings strongly encourage the exploration of additional processing alternatives (see Discussion), some of which may produce results yet superior to those here, we can provisionally recommend MSM-All alignment and the associated multimodal brain parcellation (Glasser, Coalson, et al., 2016), together with a multivariate learning model such as elastic net regression.

Finally, results for REST12 (all resting-state runs combined), using MSM-All alignment and denoising strategy A, and the multivariate learning model, are shown in **Figure 5b** together with statistical assessment using 1000 permutations. Only Openness to experience could be predicted above chance, albeit with a very small effect size (r=0.24, R^2^=0.024).

### Predicting higher-order dimensions of personality (α and β)

In previous sections, we qualitatively observed that decreasing noise in individual FC matrices by averaging data over all available resting state runs (REST12, 1 h of data) leads to improvements in prediction performance compared to session-wise predictions (REST1 and REST2, 30 min of data each). We can also decrease noise in the behavioral data, by deriving composite scores that pool over a larger number of test items than the Big Five factor scores (each factor relies solely on 12 items in the NEO-FFI). The principal component analysis presented in **Figure 2c** is a way to achieve such pooling. We therefore next attempted to predict these two principal component scores, which we refer to as α and β, from REST12 FC matrices, using denoising A and MSM-all inter-subject alignment.

α was not predicted above chance, which was somewhat expected because it loads most highly on Neuroticism, which we could not predict well in the previous section.

β was predicted above chance (p_1000_<0.002), which we also expected because it loads most highly on Openness to experience (which had r = 0.24, R^2^=0.024; **Figure 5b**). Since β effectively combines variance from Openness with that from other factors (Conscientiousness, Extraversion, and Agreeableness; see **Figure 2c**) this leads to a slight improvement in predictability, and a doubling of the explained variance (β: r = 0.27, R^2^ = 0.050). This result strongly suggests that improving the reliability of scores on the behavioral side helps boost predictability (Gignac & Bates, 2017), just as improving the reliability of FC matrices by combining REST1 and REST2 improved predictability.

## Discussion

### Summary of results

Connectome-based predictive modeling (Dubois & Adolphs, 2016; Xilin Shen et al., 2017) has been an active field of research in the past years: it consists in using functional connectivity as measured from resting-state fMRI data to predict individual differences in demographics, behavior, psychological profile, or psychiatric diagnosis. Here, we applied this approach and attempted to predict the Big Five personality factors (R. R. McCrae & Costa, 1987) from resting-state data in a large public dataset, the Human Connectome Project (N=884 after exclusion criteria). We can summarize our findings as follows.

1. We found that personality traits were not only intercorrelated with one another, but were also correlated with fluid intelligence, age, sex, handedness and other measures. We therefore regressed these possible confounds out, producing a residualized set of personality trait measures (that were, however, still intercorrelated amongst themselves).
2. Comparing different processing pipelines and data from different fMRI sessions showed generally good stability of functional connectivity across time, a prerequisite for attempting to predict a personality trait that is also stable across time.
3. We qualitatively replicated and extended a previously published finding, the prediction of a measure of fluid intelligence (Finn et al., 2015; Noble et al., 2017) from functional connectivity patterns, providing reassurance that our approach is able to predict individual differences when possible.
4. We then carried out a total of 36 different analyses for each of the five personality factors. The 36 different analyses resulted from separately analysing data from 2 sessions (establishing test-retest reliability), each with 3 different preprocessing pipelines (exploring sensitivity to how the fMRI data are processed), 2 different alignment and hard parcellation schemes (providing initial results whether multimodal surface-based alignment improves on classical volumetric alignment), and 3 different predictive models (univariate positive, univariate negative, and multivariate). Across all of these alternatives, we generally found that the MSM-All multimodal alignment together with the parcellation scheme of Glasser et al. (2016) was associated with the greatest predictability; and likewise for the multivariate model (elastic net).
5. Among the personality measures, Openness to experience showed the most reliable prediction between the two fMRI sessions, followed by Extraversion; for all other factors, predictions were often highly unstable, showing large variation depending on small changes in preprocessing, or across sessions.
6. Combining data from both fMRI sessions improved predictions. Likewise, combining behavioral data through principal component analysis improved predictions. At both the neural and behavioral ends, improving the quality of our measurements could improve predictions.
7. We best predicted the β superordinate factor, with r=0.27 and R^2^=0.05. This is highly significant as per permutation testing (though, in interpreting the statistical significance of any single finding, we note that one would have to correct for all the multiple analysis pipelines that we tested; future replications or extensions of this work would benefit from a pre-registered single approach to reduce the degrees of freedom in the analysis).

Though some of our findings achieve statistical significance in the large sample of subjects provided by the HCP, resting-state functional connectivity still only explains at most 5% of the variance in any personality score. We are thus still far from understanding the neurobiological substrates of personality (Yarkoni., 2015) (and, for that matter, of fluid intelligence which we predicted at a similar, slightly lower level; but, see (Dubois et al., 2018)). Indeed, based on this finding, it seems unlikely that findings from predictive approaches using whole-brain resting-state fMRI will inform hypotheses about specific neural systems that provide a causal mechanistic explanation of how personality is expressed in behavior.

Taken together, our approach provides important general guidelines for personality neuroscience studies using resting-state fMRI data: i) operations that are sometimes taken for granted, such as resting-state fMRI denoising (Abraham et al., 2016), make a difference to the outcome of connectome-based predictions and their test-retest reliability; ii) new inter-subject alignment procedures, such as multimodal surface matching (Robinson et al., 2014), improve performance and test-retest reliability; iii) a simple multivariate linear model may be a good alternative to the separate univariate models proposed by (Finn et al., 2015), yielding improved performance.

Our approach also draws attention to the tremendous analytical flexibility that is available in principle (Carp, 2012), and to the all-too-common practice of keeping such explorations “behind the scenes” and only reporting the “best” strategy, leading to an inflation of positive findings reported in the literature (Neuroskeptic, 2012; Simonsohn, Nelson, & Simmons, 2014). At a certain level, if all analyses conducted make sense (i.e., would pass a careful expert reviewer’s scrutiny), they should all give a similar answer to the final question (conceptually equivalent to inter-rater reliability (Dubois & Adolphs, 2016)).

#### Effect of subject alignment

The recently proposed multimodal surface matching framework uses a combination of anatomical and functional features to best align subject cortices. It improves functional inter-subject alignment over the classical approach of warping brains volumetrically (Dubois & Adolphs, 2016). For the scores that can be predicted from functional connectivity, alignment in the MSM-All space outperformed alignment in the MNI space. However, more work needs to be done to further establish the superiority of the MSM-All approach. Indeed, the parcellations used in this study differed between the MNI and MSM-All space: the parcellation in MSM-All space had more nodes (360, vs. 268) and no subcortical structures were included. Also, it is unclear how the use of resting-state data during the alignment process in the MSM-All framework interacts with resting-state based predictions, since the same data used for predictions has already been used to align subjects. Finally, it has recently been shown that the precise anatomy of each person’s brain, even after the best alignment, introduces variability that interacts with functional connectivity (Bijsterbosch et al., 2018). The complete description of brain variability at both structural and functional levels will need to be incorporated into future studies of individual differences.

#### Effect of preprocessing

We applied three separate, reasonable denoising strategies, inspired from published work (Ciric et al., 2017; Finn et al., 2015; Satterthwaite, Elliott, et al., 2013; Siegel et al., 2016) and our current understanding of resting-state fMRI confounds (Caballero-Gaudes & Reynolds, 2016; Murphy et al., 2013). The differences between the three denoising strategies in terms of the resulting test-retest reliability, based on several metrics, were not very large - yet, there were differences. Pipeline A appeared to yield the best reliability in terms of behavioral utility, while Pipeline B was best at conserving differences across subjects. Pipeline C performed worst on these metrics in our hands, despite its use of the automated artifact removal tool ICA-FIX (Salimi-Khorshidi et al., 2014); it is possible that performing CompCor and censoring are in fact detrimental after ICA-FIX (see also (Muschelli et al., 2014)). Finally, in terms of the final predictive score, all three strategies demonstrated acceptable test-retest reliability for scores that were successfully predicted.

The particular choices of pipelines that we made were intended to provide an initial survey of some commonly used schemes, but substantial future work will be needed to explore the space of possibilities more comprehensively. For instance, global signal regression - which was a part of all three chosen strategies - remains a somewhat controversial denoising step, and could be omitted if computing partial correlations, or replaced with a novel temporal ICA decomposition approach (Glasser et al., 2017). The bandpass filtering used in all our denoising approaches to reduce high frequency noise could also be replaced with alternatives such as PCA decomposition combined with “Wishart rolloff” (Glasser, Smith, et al., 2016). All of these choices impact the amount and quality of information in principle available, and how that information can be used to build a predictive model.

#### Effect of predictive algorithm

Our exploration of a multivariate model was motivated by the seemingly arbitrary decision to weight all edges equally in the univariate models (positive and negative) proposed by (Finn et al., 2015). However, we also recognize the need for simple models, given the paucity of data compared to the number of features (curse of dimensionality). We thus explored a regularized regression model that would combine information from negative and positive edges optimally, after performing the same feature-filtering step as in the univariate models. The multivariate model performed best on the scores that were predicted most reliably, yet it also seemed to have lower test-retest reliability. More work remains to be done on this front to find the best simple model that optimally combines information from all edges and can be trained in a situation with limited data.

#### Statistical significance

It is inappropriate to assess statistical significance using parametric statistics in the case of a cross-validation analysis (**Figure 4b**). However, for complex analyses, it is often the preferred option, due to the prohibitive computational resources needed to run permutation tests. Here we showed the empirical distribution of chance prediction scores for both the univariate (positive)- and multivariate-model predictions of fluid intelligence (*PMAT24_A_CR*) using denoising pipeline A in MSM-All space (**Figure 4b**). As expected, the permutation distribution is wider than the parametric estimate; it also differs significantly between the univariate and the multivariate models. This finding stresses that one needs to calculate permutation statistics for the specific analysis that one runs. The calculation of permutation statistics should be feasible given the rapid increase and ready availability of computing clusters with multiple processors. We show permutation statistics for all our key findings, but we did not correct for the multiple comparisons (5 personality factors, multiple processing pipelines). Future studies should ideally provide analyses that are pre-registered to reduce the degrees of freedom available and aid interpretation of statistical reliability.

#### Will our findings reproduce?

It is common practice in machine learning competitions to set aside a portion of your data and not look at it at all until a final analysis has been decided, and only then to run that single final analysis on the held-out data to establish out-of-sample replication. We decided not to split our dataset in that way due to its already limited sample size, and instead used a careful cross-validation framework, assessed test-retest reliability across data from different sessions, and refrained from adaptively changing parameters upon examining the final results. The current paper should now serve as the basis of a pre-registered replication, to be performed on an independent dataset (a good candidate would be the NKI enhanced dataset (Nooner et al., 2012), which also contains assessment of the Big Five).

### On the relationship between brain and personality

The best neural predictor of personality may be distinct, wholly or in part, from the actual neural mechanisms by which personality expresses itself on any given occasion. Personality may stem from a disjunctive and heterogeneous set of biological constraints that in turn influence brain function in complex ways (Yarkoni., 2015); neural predictors may simply be conceived of as “markers” of personality: any correlated measures that a machine learning algorithm could use as information, on the basis of which it could be trained in a supervised fashion to discriminate among personality traits. Our goal in this study was to find such predictions, not a causal explanation (see (Yarkoni & Westfall, 2017)). It may well someday be possible to predict personality differences from fMRI data with much greater accuracy than what we found here. However, we think it likely that, in general, such an approach will still fall short of uncovering the neural mechanisms behind personality, in the sense of explaining the proximal causal processes whereby personality is expressed in behavior on specific occasions.

### Subjective and objective measures of personality

As noted already in the introduction, it is worth keeping in mind the history of the Big Five: they derive from factor analyses of words, of the vocabularies that we use to describe people. As such, they fundamentally reflect our folk psychology, and our social inferences (“theory of mind”) about other people. This factor structure was then used to design a self-report instrument, in which participants are asked about themselves (the NEO or variations thereof). Unlike some other self-report indices (such as the MMPI), the NEO-FFI does not assess test-taking approach (e.g. consistency across items or tendency toward a particular response set), and thus, offers no insight regarding validity of any individual’s responses. This is a notable limitation, as there is substantial evidence that NEO-FFI scores may be intentionally manipulated by the subject’s response set (Furnham, 1997; Topping & O’Gorman, 1997). Even in the absence of intentional ‘faking’, NEO outcomes are likely to be influenced by an individual’s insight, impression management, and reference group effects. However, these limitations may be addressed by applying the same analysis to multiple personality measures with varying degrees of face-validity and objectivity, as well as measures that include indices of response bias. This might include ratings provided by a familiar informant, implicit-association tests (e.g. (Schnabel, Asendorpf, & Greenwald, 2008)), and spontaneous behavior (e.g. (Mehl, Gosling, & Pennebaker, 2006)). Future development of behavioral measures of personality that provide better convergent validity and discriminative specificity will be an important component of personality neuroscience.

### Limitations and Future Directions

There are several limitations of the present study that could be improved upon or extended in future work. In addition to the obvious issue of simply needing more, and/or better quality, data, there is the important issue of obtaining a better estimate of variability within a single subject. This is especially pertinent for personality traits, which are supposed to be relatively stable within an individual. Thus, collecting multiple fMRI datasets, perhaps over weeks or even years, could help to find those features in the data with the best cross-temporal stability. Indeed several such dense datasets across multiple sessions in a few subjects have already been collected, and may help guide the intelligent selection of features with the greatest temporal stability (Gordon et al., 2017; Noble et al., 2017; Poldrack et al., 2015). Against expectations, initial analyses seem to indicate that the most reliable edges in FC from such studies are not necessarily the most predictive edges (for fluid intelligence; (Noble et al., 2017)), yet more work needs to be done to further test this hypothesis. It is also possible that shorter timescale fluctuations in resting-state fMRI provide additional information (if these are stable over longer times), and it might thus be fruitful to explore dynamic FC, as some work has done (Calhoun, Miller, Pearlson, & Adalı, 2014; Jia, Hu, & Deshpande, 2014; Vidaurre, Smith, & Woolrich, 2017).

No less important would be improvements on the behavioral end, as we alluded to in the previous section. Developing additional tests of personality to provide convergent validity to the personality dimension constructs would help provide a more accurate estimate of these latent variables. Just as with the fMRI data, collecting personality scores across time should help to prioritize those items that have the greatest temporal stability and reduce measurement error.

Another limitation is signal-to-noise. It may be worth exploring fMRI data obtained while watching a movie that drives relevant brain function, rather than during rest, in order to maximize the signal variance in the fMRI signal. Similarly, it could be beneficial to include participants with a greater range of personality scores, perhaps even including those with a personality disorder. A greater range of signal both on the fMRI end and on the behavioral end would help provide greater power to detect associations.

One particularly relevant aspect of our approach is that the models we used, like most in the literature, were linear. Nonlinear models may be more appropriate, yet the difficulty in using such models is that they would require a much larger number of training samples relative to the number of features in the dataset. This could be accomplished both by accruing ever larger databases of rs-fMRI data, and by further reducing the dimensionality of the data, for instance through PCA or coarser parcellations. Alternatively, one could form a hypothesis about the shape of the function that might best predict personality scores and explicitly include this in a model.

A final important but complex issue concerns the correlation between most behavioral measures. In our analyses, we regressed out fluid intelligence, age, and sex, amongst other variables. But there are many more that are likely to be correlated with personality at some level. If one regressed out all possible measures, one would likely end up removing what one is interested in, since eventually the residual of personality would shrink to a very small range. An alternative approach is to use the raw personality scores (without any removal of confounds at all), and then selectively regress out fluid intelligence, memory task performance, mood, etc., and make comparisons between the results obtained (we provide such minimally deconfounded results in Supplementary Figure 2). This could yield insights into which other variables are driving the predictability of a personality trait. It could also suggest specific new variables to investigate in their own right. Finally, multiple regression may not be the best approach to addressing these confounds, due to noise in the measurements. Specifying confounds within a structural equation model may be a better approach (Westfall & Yarkoni, 2016).

### Recommendations for Personality Neuroscience

There are well known challenges to the reliability and reproducibility of findings in personality neuroscience, which we have already mentioned. The field shares these with any other attempt to link neuroscience data with individual differences (Dubois & Adolphs, 2016). We conclude with some specific recommendations for the field going forward, focusing on the use of resting-state fMRI data.

i. Given the effect sizes that we report here (which are by no means a robust estimate, yet do provide a basis on which to build), we think it would be fair to recommend a minimum sample size of 500 or so subjects (Schönbrodt & Perugini, 2013) for connectome-based predictions. If other metrics are used, a careful estimate of effect size that adjusts for bias in the literature should be undertaken for the specific brain measure of interest (cf. (Anderson, Kelley, & Maxwell, 2017)).
ii. A predictive framework is essential (Dubois & Adolphs, 2016; Yarkoni & Westfall, 2017), as it ensures out-of-sample reliability. Personality neuroscience studies should use proper cross-validation (in the case of the HCP, taking family structure into account), with permutation statistics. Even better, studies should include a replication sample which is held out and not examined at all until the final model has been decided from the discovery sample (advanced methods may help implement this in a more practical manner (Dwork et al., 2015)).
iii. Data sharing. If new data are collected by individual labs, it would be very important to make these available, in order to eventually accrue the largest possible sample size in a database. It has been suggested that contact information about the participants would also be valuable, so that additional measures (or retest reliability) could be collected (Mar et al., 2013). Some of these data could be collected over the internet.
iv. Complete transparency and documentation of all analyses, including sharing of all analysis scripts, so that the methods of published studies can be reproduced. Several papers give more detailed recommendations for using and reporting fMRI data, see (Dubois & Adolphs, 2016; Nichols et al., 2016; Poldrack et al., 2008). Our paper makes specific recommendation about detailed parcellation, processing and modeling pipelines; however, this is a continuously evolving field and these recommendations will likely change with future work. For personality in particular, detailed assessment for all participants, and justified exclusionary and inclusionary criteria should be provided. As suggested above, authors should consider pre-registering their study, on the Open Science Framework or a similar platform.
v. Ensure reliable and uniform behavioral estimates of personality. This is perhaps one of the largest unsolved challenges. Compared with the huge ongoing effort and continuous development of the processing and analysis of fMRI data, the measures for personality are mostly stagnant and face many problems of validity. For the time being, a simple recommendation would be to use a consistent instrument and stick with the Big Five, so as not to mix apples and oranges by using very different instruments. That said, it will be important to explore other personality measures and structures. As we noted above, there is in principle a large range of more subjective, or more objective, measures of personality. It would be a boon to the field if these were more systematically collected, explored, and possibly combined to obtain the best estimate of the latent variable of personality they are thought to measure.
vi. Last but not least, we should consider methods in addition to fMRI and species in addition to humans. To the extent that a human personality dimension appears to have a valid correlate in an animal model, it might be possible to collect large datasets, and to complement fMRI with optical imaging or other modalities. Studies in animals may also yield the most powerful tools to examine specific neural circuits, a level of causal mechanism that, as we argued above, may largely elude analyses using resting-state fMRI.

## Author contributions

J. Dubois and P. Galdi developed the overall general analysis framework and conducted some of the initial analyses for the paper. J. Dubois conducted all final analyses and produced all figures. Y. Han helped with literature search and analysis of behavioral data. L. Paul helped with literature search, analysis of behavioral data, and interpretation of the results. J. Dubois and R. Adolphs wrote the initial manuscript and all authors contributed to the final manuscript. All authors contributed to planning and discussion on this project. The authors declare no conflict of interest.

## Acknowledgments

This work was supported by NIMH grant 2P50MH094258 (RA), the Carver Mead Seed Fund (RA), and a NARSAD Young Investigator Grant from the Brain and Behavior Research Foundation (JD).

## Data Sharing

The Young Adult HCP dataset is publicly available at https://www.humanconnectome.org/study/hcp-young-adult. Analysis scripts are available in the following public repository: https://github.com/adolphslab/HCP_MRI-behavior.

---

**Supplementary Table 1.** *List of HCP subjects included in the present study.*

|  |  |  |  |  |  |  |  |  |  |  |  |  |  |  |
| --- | --- | --- | --- | --- | --- | --- | --- | --- | --- | --- | --- | --- | --- | --- |
| 100206 | 100307 | 100408 | 100610 | 101006 | 101309 | 101915 | 102311 | 102513 | 102614 | 102715 | 102816 | 103010 | 103111 | 103212 |
| 103414 | 103818 | 104012 | 104416 | 105014 | 105115 | 105216 | 105620 | 105923 | 106016 | 106319 | 106521 | 106824 | 107018 | 107321 |
| 107422 | 108020 | 108222 | 108323 | 108525 | 109325 | 110007 | 110411 | 110613 | 111009 | 111211 | 111312 | 111413 | 111514 | 111716 |
| 112112 | 112314 | 112516 | 112920 | 113215 | 113316 | 113619 | 113922 | 114217 | 114318 | 114419 | 114621 | 114823 | 115017 | 115320 |
| 115724 | 115825 | 116726 | 117021 | 117122 | 117324 | 117930 | 118023 | 118124 | 118225 | 118528 | 118730 | 118831 | 118932 | 119025 |
| 119126 | 119732 | 119833 | 120111 | 120212 | 120414 | 120515 | 120717 | 121416 | 121618 | 121921 | 122317 | 122620 | 122822 | 123824 |
| 123925 | 124422 | 124826 | 125222 | 125424 | 125525 | 126325 | 126426 | 126628 | 127226 | 127327 | 127630 | 127731 | 127832 | 127933 |
| 128026 | 128127 | 128632 | 128935 | 129028 | 129129 | 129331 | 129634 | 130013 | 130114 | 130316 | 130417 | 130619 | 130720 | 130821 |
| 130922 | 131217 | 131419 | 131722 | 131823 | 132017 | 132118 | 133019 | 133625 | 133827 | 133928 | 134021 | 134223 | 134324 | 134425 |
| 134627 | 134728 | 134829 | 135124 | 135225 | 135528 | 135629 | 135730 | 135932 | 136126 | 136227 | 136631 | 136833 | 137027 | 137128 |
| 137229 | 137532 | 137633 | 137936 | 138130 | 138231 | 138332 | 138534 | 138837 | 139233 | 139637 | 139839 | 140117 | 140420 | 141422 |
| 142828 | 143224 | 143325 | 143426 | 143830 | 144125 | 144226 | 144428 | 144832 | 144933 | 145127 | 145632 | 145834 | 146129 | 146331 |
| 146432 | 146533 | 146735 | 146836 | 146937 | 147636 | 147737 | 148032 | 148133 | 148335 | 148436 | 148840 | 148941 | 149236 | 149337 |
| 149539 | 149741 | 149842 | 150423 | 150625 | 150726 | 150928 | 151425 | 151627 | 151829 | 151930 | 152225 | 152427 | 152831 | 153025 |
| 153126 | 153227 | 153429 | 153631 | 153732 | 153934 | 154229 | 154431 | 154532 | 154734 | 154835 | 154936 | 155635 | 155938 | 156031 |
| 156334 | 156435 | 156536 | 157336 | 157437 | 157942 | 158035 | 158136 | 158338 | 158540 | 158843 | 159239 | 159340 | 159441 | 159744 |
| 159946 | 160123 | 160729 | 161630 | 161731 | 162026 | 162228 | 162329 | 162733 | 162935 | 163129 | 163432 | 164030 | 164636 | 164939 |
| 165032 | 165436 | 165638 | 165840 | 165941 | 166438 | 167036 | 167238 | 167440 | 167743 | 168139 | 168341 | 168745 | 168947 | 169343 |
| 169444 | 169545 | 169747 | 169949 | 170631 | 171330 | 171633 | 172029 | 172130 | 172332 | 172433 | 172534 | 172938 | 173334 | 173435 |
| 173536 | 173637 | 173738 | 173839 | 173940 | 174437 | 174841 | 175136 | 175237 | 175338 | 175540 | 175742 | 176037 | 176441 | 176542 |
| 176744 | 176845 | 177140 | 177241 | 177645 | 177746 | 178142 | 178243 | 178748 | 178849 | 178950 | 179245 | 179346 | 180230 | 180432 |
| 180533 | 180735 | 180836 | 180937 | 181131 | 181232 | 182032 | 182739 | 182840 | 183034 | 185038 | 185139 | 185341 | 185442 | 185846 |
| 185947 | 186040 | 186141 | 186444 | 186545 | 186848 | 187143 | 187345 | 187547 | 187850 | 188145 | 188347 | 188448 | 188549 | 189349 |
| 189450 | 189652 | 190031 | 191033 | 191235 | 191336 | 191841 | 191942 | 192035 | 192237 | 192439 | 192540 | 192641 | 192843 | 193239 |
| 193845 | 194140 | 194443 | 194645 | 194746 | 194847 | 195041 | 195445 | 195647 | 195849 | 195950 | 196144 | 196346 | 196750 | 196952 |
| 197550 | 198047 | 198350 | 198451 | 198653 | 198855 | 199251 | 199352 | 199453 | 199655 | 199958 | 200008 | 200109 | 200311 | 200513 |
| 200614 | 200917 | 201111 | 201414 | 201515 | 201818 | 202113 | 202719 | 202820 | 203418 | 203923 | 204016 | 204218 | 204319 | 204420 |
| 204521 | 204622 | 205119 | 205725 | 205826 | 206222 | 206323 | 206525 | 206727 | 206828 | 206929 | 208024 | 208125 | 208226 | 208327 |
| 208630 | 209127 | 209228 | 209329 | 209935 | 210011 | 210112 | 210617 | 211114 | 211215 | 211316 | 211417 | 211619 | 211821 | 211922 |
| 212015 | 212318 | 212419 | 212823 | 213017 | 213421 | 213522 | 214019 | 214221 | 214423 | 214524 | 214625 | 214726 | 217126 | 217429 |
| 219231 | 220721 | 221319 | 224022 | 227533 | 228434 | 231928 | 237334 | 238033 | 239136 | 239944 | 245333 | 246133 | 248339 | 249947 |
| 250932 | 251833 | 255639 | 255740 | 256540 | 257542 | 257845 | 257946 | 263436 | 268749 | 268850 | 270332 | 274542 | 275645 | 280739 |
| 280941 | 281135 | 283543 | 285345 | 285446 | 286347 | 286650 | 287248 | 289555 | 290136 | 293748 | 297655 | 298455 | 299154 | 299760 |
| 300618 | 300719 | 303119 | 303624 | 304020 | 304727 | 305830 | 307127 | 308129 | 309636 | 310621 | 314225 | 316633 | 316835 | 317332 |
| 318637 | 320826 | 321323 | 322224 | 325129 | 329440 | 329844 | 330324 | 333330 | 334635 | 336841 | 339847 | 341834 | 342129 | 346945 |
| 348545 | 349244 | 350330 | 352738 | 353740 | 356948 | 358144 | 360030 | 361234 | 361941 | 365343 | 366042 | 368551 | 368753 | 371843 |
| 376247 | 377451 | 378857 | 379657 | 380036 | 381038 | 381543 | 385046 | 385450 | 386250 | 387959 | 389357 | 391748 | 392750 | 393550 |
| 394956 | 395251 | 395756 | 397154 | 397760 | 401422 | 406432 | 406836 | 412528 | 413934 | 414229 | 415837 | 419239 | 421226 | 422632 |
| 424939 | 429040 | 432332 | 436239 | 436845 | 441939 | 445543 | 448347 | 449753 | 453441 | 454140 | 456346 | 459453 | 461743 | 463040 |
| 467351 | 469961 | 475855 | 479762 | 480141 | 481042 | 481951 | 485757 | 486759 | 497865 | 499566 | 500222 | 506234 | 510326 | 512835 |
| 513130 | 513736 | 516742 | 517239 | 518746 | 519647 | 519950 | 520228 | 522434 | 523032 | 524135 | 525541 | 529549 | 529953 | 530635 |
| 531536 | 531940 | 536647 | 540436 | 541640 | 541943 | 545345 | 547046 | 548250 | 552241 | 552544 | 553344 | 555651 | 555954 | 557857 |
| 558657 | 558960 | 559053 | 559457 | 561242 | 561444 | 561949 | 562446 | 565452 | 566454 | 567052 | 567759 | 567961 | 568963 | 570243 |
| 571144 | 572045 | 573249 | 573451 | 578057 | 579665 | 579867 | 580044 | 580347 | 580650 | 580751 | 581450 | 583858 | 585256 | 585862 |
| 586460 | 587664 | 588565 | 589567 | 590047 | 592455 | 594156 | 597869 | 598568 | 599065 | 599469 | 599671 | 601127 | 604537 | 611938 |
| 613538 | 615441 | 615744 | 616645 | 617748 | 618952 | 620434 | 622236 | 623844 | 626648 | 627852 | 628248 | 634748 | 635245 | 638049 |
| 644246 | 645450 | 645551 | 647858 | 654350 | 654552 | 654754 | 656253 | 656657 | 657659 | 660951 | 663755 | 664757 | 665254 | 667056 |
| 668361 | 671855 | 672756 | 673455 | 675661 | 677766 | 679568 | 679770 | 680452 | 683256 | 685058 | 687163 | 690152 | 692964 | 693764 |
| 694362 | 695768 | 700634 | 702133 | 704238 | 705341 | 707749 | 709551 | 715041 | 715647 | 715950 | 720337 | 723141 | 724446 | 725751 |
| 727553 | 727654 | 728454 | 729254 | 729557 | 731140 | 732243 | 734045 | 737960 | 742549 | 744553 | 748258 | 749058 | 749361 | 751348 |
| 751550 | 753150 | 753251 | 756055 | 757764 | 759869 | 761957 | 763557 | 765056 | 765864 | 767464 | 769064 | 770352 | 771354 | 773257 |
| 774663 | 782561 | 783462 | 784565 | 788674 | 789373 | 792564 | 792766 | 793465 | 800941 | 802844 | 803240 | 809252 | 812746 | 814548 |
| 814649 | 815247 | 816653 | 818455 | 818859 | 820745 | 825048 | 825553 | 825654 | 826353 | 826454 | 827052 | 828862 | 832651 | 833148 |
| 835657 | 837560 | 841349 | 843151 | 844961 | 845458 | 849264 | 849971 | 852455 | 856766 | 857263 | 859671 | 861456 | 865363 | 867468 |
| 870861 | 871762 | 871964 | 872158 | 872562 | 872764 | 873968 | 877168 | 877269 | 878776 | 878877 | 880157 | 882161 | 884064 | 885975 |
| 886674 | 887373 | 888678 | 889579 | 891667 | 894067 | 894673 | 894774 | 896778 | 896879 | 898176 | 899885 | 901139 | 901442 | 902242 |
| 904044 | 905147 | 907656 | 908860 | 910241 | 910443 | 911849 | 912447 | 917255 | 917558 | 923755 | 926862 | 927359 | 930449 | 932554 |
| 933253 | 942658 | 943862 | 951457 | 952863 | 955465 | 957974 | 958976 | 959574 | 962058 | 965367 | 966975 | 969476 | 970764 | 971160 |
| 978578 | 979984 | 983773 | 984472 | 987074 | 987983 | 989987 | 990366 | 991267 | 992673 | 992774 | 993675 | 994273 | 996782 |  |

**Supplementary Figure 1.**
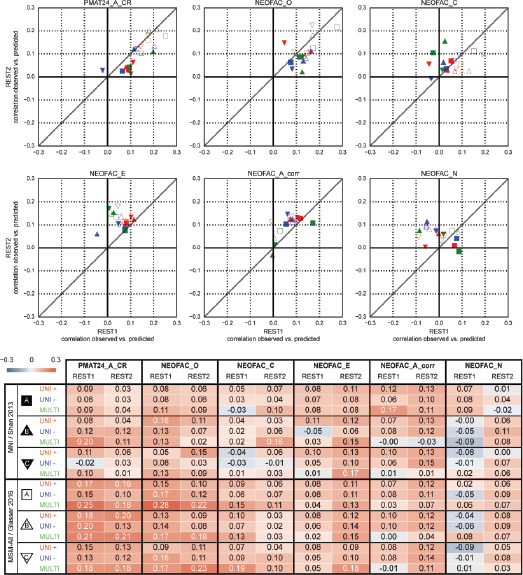
Test-retest prediction results with minimal deconfounding. Only variables that are unlikely to be causally related to personality are regressed out of the scores: brain size, motion, and MB recon version.

